# A barley gene cluster for the biosynthesis of diterpenoid phytoalexins

**DOI:** 10.1101/2021.05.21.445084

**Authors:** Yaming Liu, Gerd U. Balcke, Andrea Porzel, Lisa Mahdi, Anja Scherr-Henning, Ulschan Bathe, Alga Zuccaro, Alain Tissier

## Abstract

Phytoalexins are specialized metabolites that are induced upon pathogen infection and contribute to the defense arsenal of plants. Maize and rice produce multiple diterpenoid phytoalexins and there is evidence from genomic sequences that other monocots may also produce diterpenoid phytoalexins. Here we report on the identification and characterization of a gene cluster in barley (*Hordeum vulgare* cv. Golden Promise) that is involved in the production of a set of labdane-related diterpenoids upon infection of roots by the fungal pathogen *Bipolaris sorokiniana*. The cluster is localized on chromosome 2, covers over 600 kb and comprises genes coding for a (+)-copalyl diphosphate synthase (HvCPS2), a kaurene synthase like (HvKSL4) and several cytochrome P450 oxygenases (CYPs). Expression of HvCPS2 and HvKSL4 in yeast and *Nicotiana benthamiana* resulted in the production of a single major product, whose structure was determined to be of the cleistanthane type and was named hordediene. Co-expression of HvCPS2, HvKSL4 and one of the CYPs from the cluster (CYP89E31) afforded two additional products, hordetriene and 11-hydroxy-hordetriene. Both of these compounds could be detected in extracts of barley roots infected by *B. sorokiniana*, validating the function of these genes *in planta*. Furthermore, diterpenoids with multiple oxidations and with molecular masses of 316, 318 and 332 were induced in infected barley roots and secreted in the medium, indicating that additional oxidases, possibly from the same genomic cluster are involved in the production of these phytoalexins. Our results provide the basis for further investigation of the role of this gene cluster in the defense of barley against pathogens and more generally in the interaction with the microbiome.

## Introduction

Monocotyledons contribute some of the most important staple crops worldwide, including the three major ones maize (*Zea mays*), wheat (*Triticum aestivum*) and rice (*Oryza sativa*) that cover a large part of calorie intake by humans worldwide (Awika, 2011). Behind these three species, barley (*Hordeum vulgare*) is the fourth most important grain crop with an annual production of over 140 million tons and with an harvest area of almost 48 million ha worldwide in 2018 (**Table S1**). Barley is grown in temperate climates and primarily for animal feed, but also to provide substrate for the fermentation of beverages (e.g. beer), and for a range of health promoting products. It is one of the earliest cultivated crops with supporting archaeological evidence from the fertile crescent region dating to as far back as over 10,000 years before present (BP) (Badr et al., 2000). The initial use of barley was for food, but it was then later replaced by wheat (Riehl, 2019). Cultivated barley is one of 31 *Hordeum* species, with *H. vulgare* subsp. *spontaneum* believed to be the wild ancestor of cultivated barley (Badr et al., 2000). The close relatedness of barley to wheat and its diploid nature (2n = 14) make it a relevant model species for the study of temperate cereal crops. A first draft of its genome was published in 2016 and was recently updated (Beier et al., 2017; Mascher et al., 2017; Monat et al., 2019). Furthermore, genetic transformation and gene editing in barley are now well established, providing a complete toolbox for functional genetics in this species (Hensel, 2020).

Plants synthesize a complex array of secondary metabolites that contribute to the response and adaptation to a range of biotic or abiotic stresses. These metabolites can be produced constitutively, in a tissue specific manner or upon challenge by specific stresses, be they biotic or abiotic. Whereas some metabolites are common to a wide range of species, others are restricted to a species or to a taxon, thereby determining a species or taxon metabolite signature, a feature that led to the denomination “specialized metabolites” (Pichersky et al., 2006; Pichersky and Lewinsohn, 2011).

Plant pathogenic fungi impose a major burden on crop yield, and this impact is expected to increase with climate change (Miedaner and Juroszek, 2021). As the use of agrochemicals is increasingly under scrutiny by environmental agencies and while single gene-for-gene resistance may be rapidly overcome in a changing climate, there is a strong need for more durable resistance traits. The fungus *Bipolaris sorokiniana* (syn. *Cochliobolus sativus*) is the pathogenic agent of spot blotch and root rot in wheat and barley and is particularly prevalent in regions with a warmer climate (Rosyara et al., 2010) and therefore represents a typical future important threat in regions with a temperate climate in the context of global warming. *B. sorokiniana* can infect both aerial and underground parts of the plant but knowledge on how it interacts with roots is still rather limited (Sarkar et al., 2019).

There is widespread evidence that infection of plants by microbial pathogens triggers the production of a secondary metabolites (also called specialized metabolites) that exhibit antimicrobial or antioxidant activities (Ahuja et al., 2012). These compounds, called phytoalexins, are not restricted to a particular chemical class, with examples among phenylpropanoids, alkaloids or terpenoids (Ahuja et al., 2012). This is also the case in Monocotyledons, a plant clade that includes some of the most important food crops worldwide. In barley, there is a number of reports of various phytoalexins produced in response to diverse pathogens. These include phenylamides, such as the dimeric hordatine A and B, the indole-derived gramine, benzoxazinones such as 2,4-dihydroxy-1,4-benzoxazin-3-one (DIBOA), methoxychalcones as well as tyramine and related amines (Ishihara et al., 2017; Ube et al., 2017; Ube et al., 2021). However, in contrast to other monocotyledon crops such as maize and rice, no sesqui- or diterpenoid phytoalexins have been identified in barley yet. There is now extensive data available on the nature and biosynthesis of a range of terpenoid phytoalexins in these important crop species. Rice produces several classes of labdane-related diterpenoids, including momilactones (A and B) (Kato et al., 1973; Cartwright et al., 1977), phytocassanes, oryzalexins (Akatsuka et al., 1983; Kono et al., 1984; Sekido et al., 1986; Kato et al., 1993, 1994) and oryzalides (Watanabe et al., 1990; Kono et al., 1991) as well as the macrocyclic *ent*-oxodepressin (Inoue et al., 2013). Maize produces dolabralexins and kauralexins, both labdane-related diterpenoid phytoalexins (Schmelz et al., 2011), as well as zealexins, which are sesquiterpenoids (Huffaker et al., 2011).

Biosynthesis of terpenoids starts by the conversion of linear isoprenyl diphosphate chains by terpene synthases to either linear or cyclic terpenes or terpene alcohols. In the case of diterpenoids, the precursor is typically all-*trans*-geranylgeranyl diphosphate (here abbreviated as GGPP) (Bohlmann et al., 1998), although in some isolated cases it is the all-*cis*-isomer, nerylneryl diphoshate (Zi et al., 2014). Diterpene synthases (diTPS) are classified according to the mechanism underlying the initiation of the cyclization reaction. Thus class I diTPS initiate the reaction by dephosphorylation whereas class II do it by protonation of the terminal isoprenic double bond (Peters, 2010). The main products of class II diTPS are *ent*-copalyl diphosphate (*ent*-CPP), the precursor of the gibberellins, and the other stereoisomers *syn*-copalyl diphosphate (*syn*-CPP) and CPP of normal configuration (Peters, 2010). In addition, there are other products of class II diTPS with a slightly different core structure, such as clerodienyl or halimadienyl diphosphates, as well as products with an alcohol function (Nakano et al., 2005; Sallaud et al., 2012; Pelot et al., 2017). Because the products of class II diTPS still contain a diphosphate group, class I diTPS can convert them to olefinic diterpenes or diterpene alcohols. The sequential reactions catalysed by class II and class I diTPS lead to the broad group of labdane-related diterpenes, which have in common a core bicyclic decalin ring structure (Peters, 2010). Apart from *ent*-oxodepressin, which has a macrocyclic structure, all diterpenoid phytoalexins from monocots identified so far belong to the labdane-related group. Following cyclization by diTPS, diterpene backbones are then oxidized at different positions and in a stereospecific way. Cytochrome P450 oxygenases (CYPs) are the most frequently involved in these oxidations (Bathe and Tissier, 2019), but other classes of enzymes such as 2-oxoglutarate dependent dioxygenases in gibberellin biosynthesis (Hedden and Kamiya, 1997) or short-chain dehydrogenases/reductases as in momilactone biosynthesis (Kitaoka et al., 2016) can also play a role in functionalizing diterpenes. These oxidations can sometimes lead to backbone rearrangements and, importantly, provide anchoring points, such as hydroxyl or carboxyl groups, for further modifications by conjugating enzymes (Long et al., 2008; Rontein et al., 2008). Thus, sugar, acyl, or benzoyl groups can decorate the oxidized diterpene core and provide added functionalities.

In recent years the elucidation of the biosynthesis of rice and maize diterpenoid phytoalexins has progressed significantly (Murphy and Zerbe, 2020). In rice, the labdane-related diterpenoids are derived from either *ent*- or *syn*-copalyl diphosphate, whereas the maize diterpenoid phytolexins derive from *ent*-CPP. In addition to the diTPS, which produce the diterpene backbones of these phytoalexins, a number of CYPs are involved in the functionalization of these backbones. Notably, both in maize and rice, some of the genes for the biosynthesis of terpenoid phytoalexins occur are physically associated in chromosomal clusters (Shimura et al., 2007; Wang et al., 2011; Ding et al., 2020; Liang et al., 2021).

Here we present the identification and characterization of a diterpenoid phytoalexin biosynthesis cluster in barley. We show that genes in this cluster are strongly induced in roots by a barley fungal pathogen, *B. sorokiniana* and we characterize the first biosynthesis steps, consisting of a copalyl diphosphate synthase, a kaurene synthase like and a CYP. Notably, the diterpene backbone produced by the CPS/KSL enzymes belong to the cleistanthane group of labdane-related diterpenoids. To the best of our knowledge, this diterpene backbone has not been detected yet in grasses. We also detected diterpenoid phytoalexins in barley roots and in high amounts in the root exudate as well as the intermediates produced by the first three enzymes of the pathway.

## Results

### Identification of a diterpenoid biosynthesis gene cluster in barley chromosome 2

In previous work, we generated transcriptome data of barley roots infected with either a fungal pathogen, *Bipolaris sorokiniana*, or a beneficial root endophyte, *Serendipita vermifera*, or both in local or systemic context (Sarkar et al., 2019). We observed that a number of genes from the MEP pathway as well as a number of genes from a region in chromosome 2 encoding terpene synthases and cytochrome P450 oxygenases are strongly induced by the fungal pathogen *B. sorokiniana* and moderately by *S. vermifera* (**Fig. 1**). In the MorexV2 version of the barley genome (Monat et al., 2019), this cluster spans over 600 kb and contains one gene encoding a copalyl diphosphate synthase (CPS), one kaurene-synthase like (KSL), 8 CYP and one asparaginase (**Fig. 1**). The cluster also harbors a number of pseudogenes, including five CYPs and four CPS. As is frequently the case in regions of the genome with multiple duplications, the early versions of the genome sequence (including MorexV1 and MorexV2) still contain a number of gaps, and it is likely that the number of genes and pseudogenes in this cluster evolves as more accurate sequences become available.

**Figure 1.**
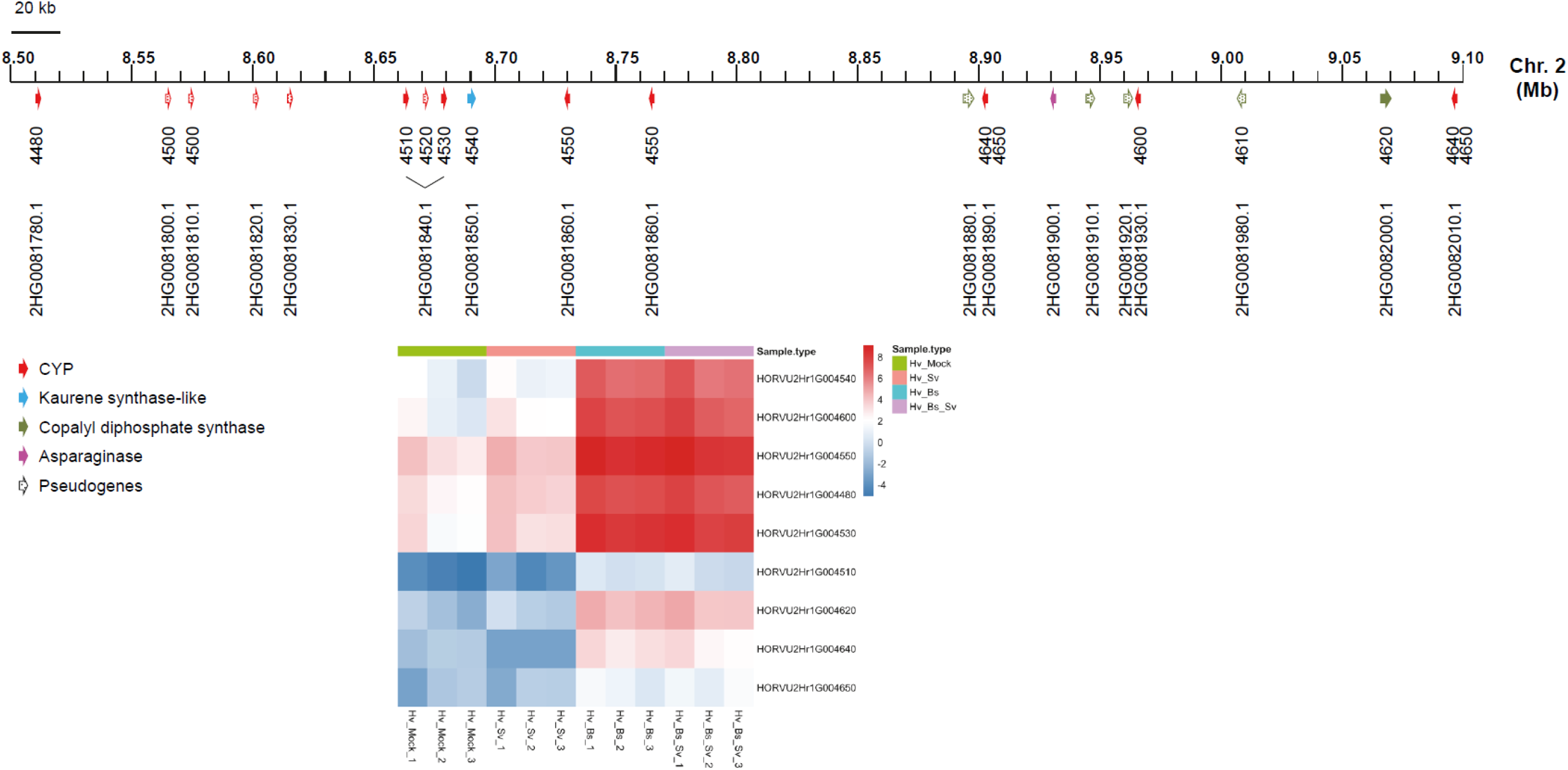
Overview of the barley chromosome 2 cluster for diterpenoid biosynthesis. The scale on top indicates the position on chromosome 2 based on the latest annotation of the barley genome (Monat et al., 2019). The genes are represented by arrows underneath the scale. The color code indicates to which gene family they belong and the dotted pattern the presence of a pseudogene. The first row of numbers below the scale indicates the last 4 digits of the gene ID according to MorexV1 gene models and the second row gives the gene identification in the MorexV2 version of the barley genome. The colored bars in the lower part represent gene expression values as log2-transformed FKPM values for genes that show differential gene expression (|fold change| > 2, data from Sarkar et al., 2019). Each square for a sample type represents data from a biological replicate. The samples types are indicated by the following color code: purple: barley root co-inoculated with *B. sorokiniana* and *S. vermifera*; turquoise blue: barley roots inoculated with *B. sorokiniana*; pink: barley roots inoculated with S. vermifera; green: barley root mock inoculated.

### Phylogenetic analysis of the CPS and KSL genes in the chromosome 2 cluster

To better characterize the genes of the cluster, we performed a phylogenetic analysis. Earlier publications reported on the identification and biochemical characterization of HvCPS1 as an *ent*-CDP synthase (Spielmeyer et al., 2004; Wu et al., 2012). The strong similarity of HvCPS1 to TaCPS3 and TaCPS4, both of which are *ent*-CDP synthases, and the exclusive presence of *ent*-CDP synthases in the same branch underscores the distinct evolutionary conservation of *ent*-CDP synthases in monocots (**Fig. 2**). The CPS in the chromosome 2 cluster has not been characterized yet and we propose to name it HvCPS2. HvCPS2 shares high similarity to TaCPS2, a (+)-CDP synthase, and belongs to the same branch as OsCPS4, a *syn*-CDP synthase, and TaCPS1, an *ent*-CDP synthase. Thus, although these data support a role of HvCPS2 in specialized diterpenoid metabolism, they do not allow us to predict its actual biochemical activity with high confidence.

**Figure 2.**
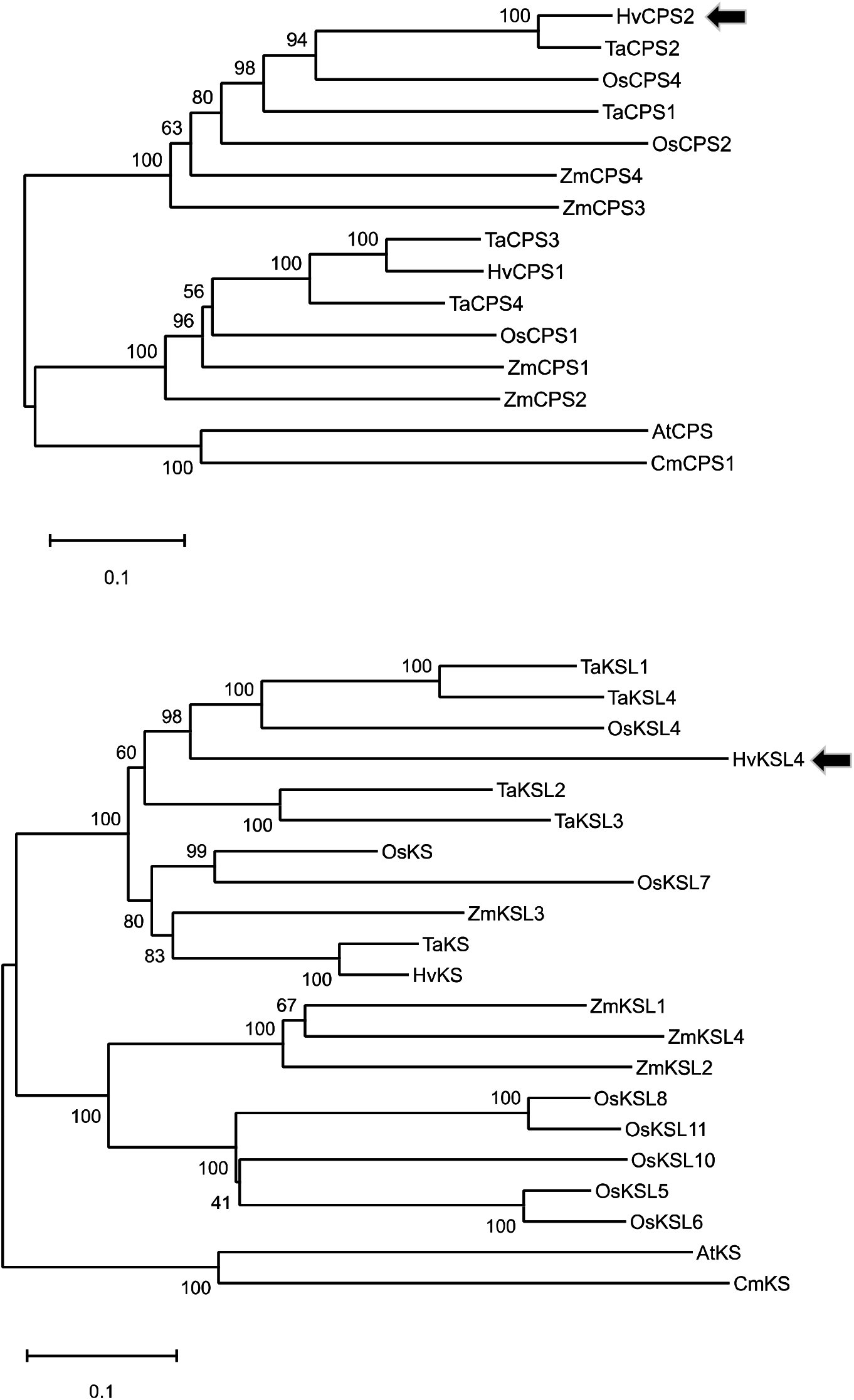
Phylogenetic analysis of HvCPS2 and HvKSL4. The sequences indicated were aligned and processed with the MEGA X software (Kumar et al., 2018) with the maximum likelihood method and 1000 bootstrap replications. The consensus trees are shown with the bootstrap values for the individual branches shown. The HvCPS2 and HvKSL4 sequences are indicated by grey arrows. The list of sequences used is provided in **Tables S2 and S3**.

The kaurene synthase-like of the chromosome 2 cluster was already identified previously and named HvKSL4 (Li et al., 2016). In the phylogenetic tree comprising other monocot KSL enzymes (**Fig. 2**), it is in the same branch as TaKSL1, TaKSL4 and OsKSL4. The substrate of TaKSL1 and TaKSL4 is (+)-CDP (Zhou et al., 2012), while that of OsKSL4 is *syn*-CDP (Otomo et al., 2004). In the neighboring branch are TaKSL2 and TaKSL3. The function of TaKSL3 is unknown but TaKSL2 also uses (+)-CDP as substrate (Zhou et al., 2012). The clear separation from KSL enzymes that use *ent*-CDP as substrate, including the barley *ent*-kaurene synthase (HvKS), indicates that HvKSL4 most likely uses either (+)-CDP or *syn*-CDP as substrate.

### HvCPS2 is a (+)-CDP synthase and HvKSL4 produces a cleistanthane-type backbone

To determine the biochemical activity of HvCPS2, we expressed it in yeast using our Golden Gate yeast cloning system (Scheler et al., 2016) together with RoMiS, the miltiradiene synthase from rosemary (*Rosmarinus officinalis*) (Brückner et al., 2014), with CcKS, an *ent*-kaurene synthase from coffee (*Coffea canephora*), or with HvKSL4. Co-expression with RoMiS yielded the expected diterpene product (miltiradiene), whereas no *ent*-kaurene could be detected with CcKS, demonstrating that HvCPS2 produces (+)-CDP **(Fig. 3**). Co-expression of HvCPS2 and HvKSL4 in yeast yielded a novel product with a molecular ion of 272 (**Fig. 3**), indicating that it is a diterpene olefin. Besides this major product, several much more minor products could also be detected. When expressing truncated versions of HvCPS2 and HvKSL4 together with a cytosolic geranylgeranyl diphosphate synthase (GGPPS) and a truncated version of hydroxymethylglutaryl CoA-reductase (tHMGR) in *Nicotiana benthamiana*, the same product could be detected (**Fig. S1**). Since there was no significant match in the NIST database (Mass Spectrometry Data Center, http://chemdata.nist.gov), we purified the main product and determined its structure by nuclear magnetic resonance (NMR) spectroscopy (see **Table S4**). The product was determined to have a cleistanthane backbone, with two double bonds in the C-ring at positions C8-C9 and C12-C13, a methyl group attached to C13 and an ethyl group attached to C14 in the α-configuration (**Fig. 3**). Cleistanthanes constitute a relatively small group of diterpenes that occur in some plants and fungi [see for example: (Kaufman et al., 1987; Riehl and Pinto, 2000; Shiono et al., 2010; Zheng et al., 2018)]. We could not find published reports of the exact same structure and we therefore named it hordediene (alternatively: cleistantha-8(9),12(13)-diene).

**Figure 3.**
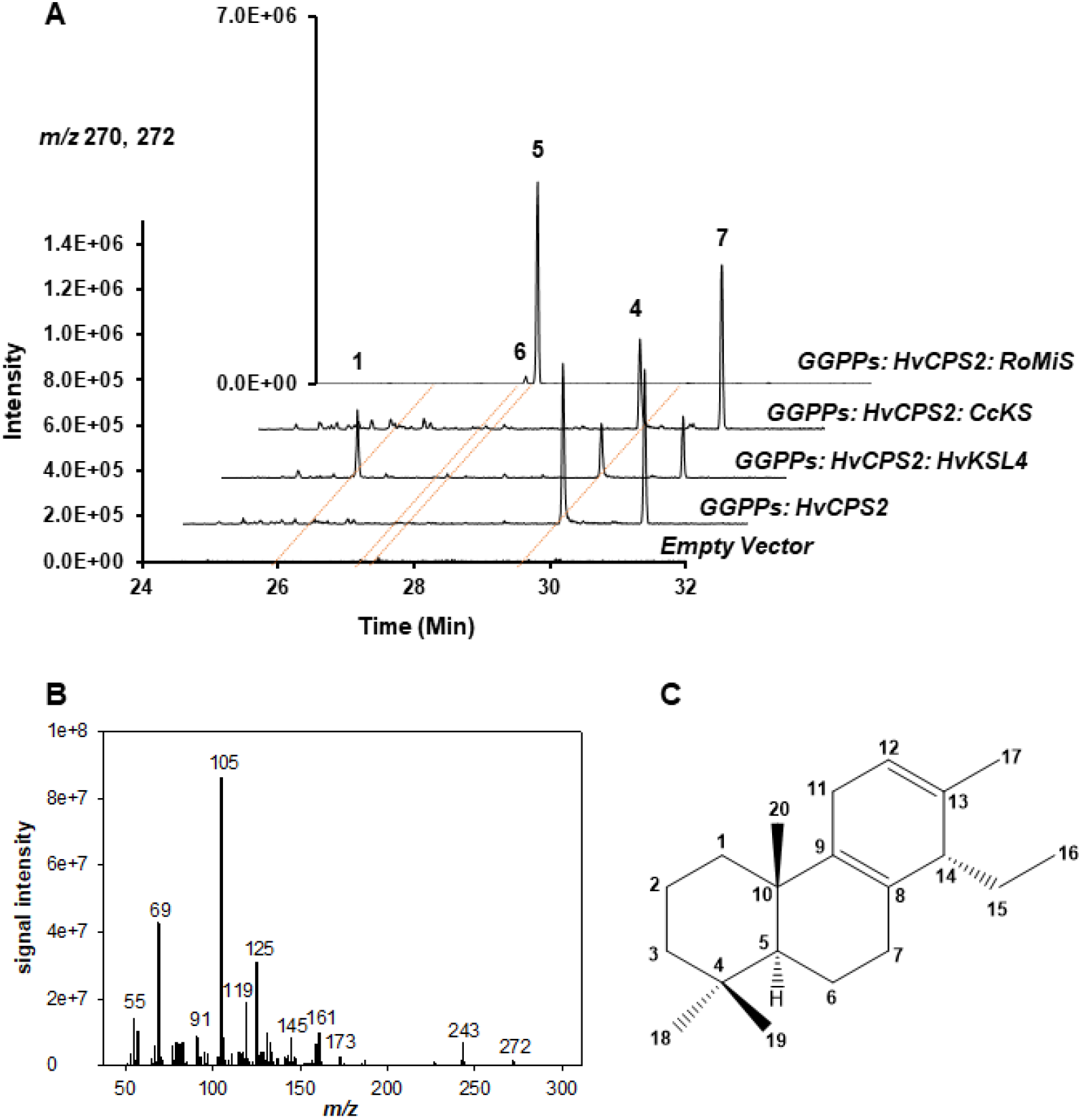
Expression of HvCPS2 and HvKSL4 in yeast. A. Selected ion (*m/z* 270 and 272) GC-MS chromatograms of *n*-hexane extracts of yeast strains expressing the gene combinations indicated on the right. CcKS: *ent*-kaurene synthase from coffee (*Coffea canephora*); RoMiS, miltiradiene synthase from rosemary (*Rosmarinus officinalis*). **1**: hordediene; **4:** (+)-copalol; **5**: miltiradiene; **6**: abietatriene; **7**: unknown copalol derivative. B. EI mass spectrum of hordediene **(1)**. Structure of hordediene as determined by NMR.

### Phylogenetic analysis of the CYPs in the chromosome 2 cluster

Of the eight CYP-encoding genes in the chromosome 2 cluster, only five are expressed at significant levels and display an expression pattern similar to that of *HvCPS2* and *HvKSL4*. Furthermore in the latest annotation of the barley genome, two of these genes merged into a single one so that only four CYP-encoding genes can be considered in the current version of the barley genome (**Table 1**). A phylogenetic analysis of these sequences shows that three of them are most similar to OsCYP99A2 and to OsCYP99A3 from rice (see **Fig. S2 and Table S5)**. Although no biochemical function could be determined for OsCYP99A2 yet, OsCYP99A3 plays a role in the biosynthesis of momilactones by oxidizing *syn*-labdane related diterpenes – *syn*-pimaradiene and *syn*-stemodene – at the C19 position to generate a carboxylic acid function (Wang et al., 2011). Interestingly, OsCYP99A2 and OsCYP99A3 are located in the same tandem cluster next to OsCPS4 and OsKSL4, a situation highly reminiscent of the barley cluster presented here (Shimura et al., 2007).

**Table 1.** List of genes from the chromosome 2 diterpenoid phytoalexin cluster.

| Genome version |  |  | Gene status | Transcript ID | Protein ID | Name |
| --- | --- | --- | --- | --- | --- | --- |
| MorexV1 (2017) | MorexV2 (2019) | MorexV3 (2021) |  |  |  |  |
| 2Hr1G004480 | 2HG0081780 | 2HG0099280 | FL | AK252527.1 |  | CYP99A66 |
| 2Hr1G004510<br>2Hr1G004520 | 2HG0081840 | 2HG0099340 | Pseudo |  |  |  |
| 2Hr1G004530 | 2HG0081840 | 2HG0099350 | FL |  | KAE8771735 | CYP99A67 |
| 2Hr1G004540 | 2HG0081850 | 2HG0099360 | FL | AK370792 |  | HvKSL4 |
| 2Hr1G004550 | 2HG0081860<br>UnG0627890<br>UnG0631950 | 2HG0099370<sup>a</sup><br>2HG0099420<sup>a</sup><br>2HG0099430<sup>a</sup> | FL | AK369243 |  | CYP89E31 |
| NP | 2HG0081880 | 2HG0099470 | Pseudo |  |  | ψCPS1 |
| 2Hr1G004600 | 2HG0081930 | 2HG0099550 | FL |  |  | CYP99A68<sup>b</sup> |
| 2Hr1G004610 | 2HG0081980 | 2HG0099550 | FL |  |  | CYP99A68<sup>b</sup> |
| 2Hr1G004640<br>2Hr1G004650 | 2HG0081890<br>2HG0082010<br>2HG0081930<br>UnG0636400 | 2HG0099480 | FL |  |  | CYP99A68<sup>b</sup> |
|  | 2HG0081920 | 2HG0099500 | Pseudo |  |  | ψCPS2 |
| 2Hr1G004620 | 2HG0082000 | 2HG0099570 | FL | AK364238 |  | HvCPS2 |

Notes: (a) the amino acid sequences of these two genes are identical. The sequences only differ in the 5'-UTR. (b) the amino acid sequences are identical except for one amino acid change in 2Hr1G004600

The fourth CYP shares strongest similarity to enzymes of the CYP89 clan, with the closest homolog in *Arabidopsis thaliana* being CYP89A2 (AT1G64900), whose biochemical function is unknown yet. The most closely related protein sequences in the databases are homologs from other crops, e.g. *Triticum aestivum* or *Setaria viridis*, but these are of unknown function as well. The assigned identification for this CYP is CYP89E31.

### Characterization of HvCYP89E31

We first focused on the characterization of HvCYP89E31 by expressing it in yeast. We cloned a yeast optimized sequence in a vector together with HvCPS2 and HvKSL4 and in addition with the cytochrome P450 reductase from Arabidopsis. Analysis of the extracts by gas chromatography couple to electron ionization mass spectrometry (GC/EI-MS) showed the presence of two additional peaks, with a molecular *m/z* of 270 (product **2**) and 286 (product **3**), respectively (**Fig. 4**). These masses indicate one and two additional degrees of oxidation. Next, we also isolated microsomes from a yeast strain expressing CYP89Ax and performed in vitro assays with hordediene as a substrate. GC-MS analysis of the extracts from this reaction showed the same products that are produced by yeast expressing all three genes (**Fig. 4**). This conclusively demonstrates that the new products detected are directly derived from hordediene. We also expressed those three genes in *N. benthamiana* as described above for HvCPS2 and HvKSL4 and found the exact same products present in hexane extracts (**Fig. S1**). The products were purified from a large scale yeast culture and their structure determined by NMR (**Tables S6 and S7**). Compound **2** has an aromatized C-ring whereas compound **3** has a hydroxyl group on position 11 (**Fig. 4**). We propose to name these compounds hordetriene (**2**) and 11-hydroxy-hordetriene (**3**).

**Figure 4.**
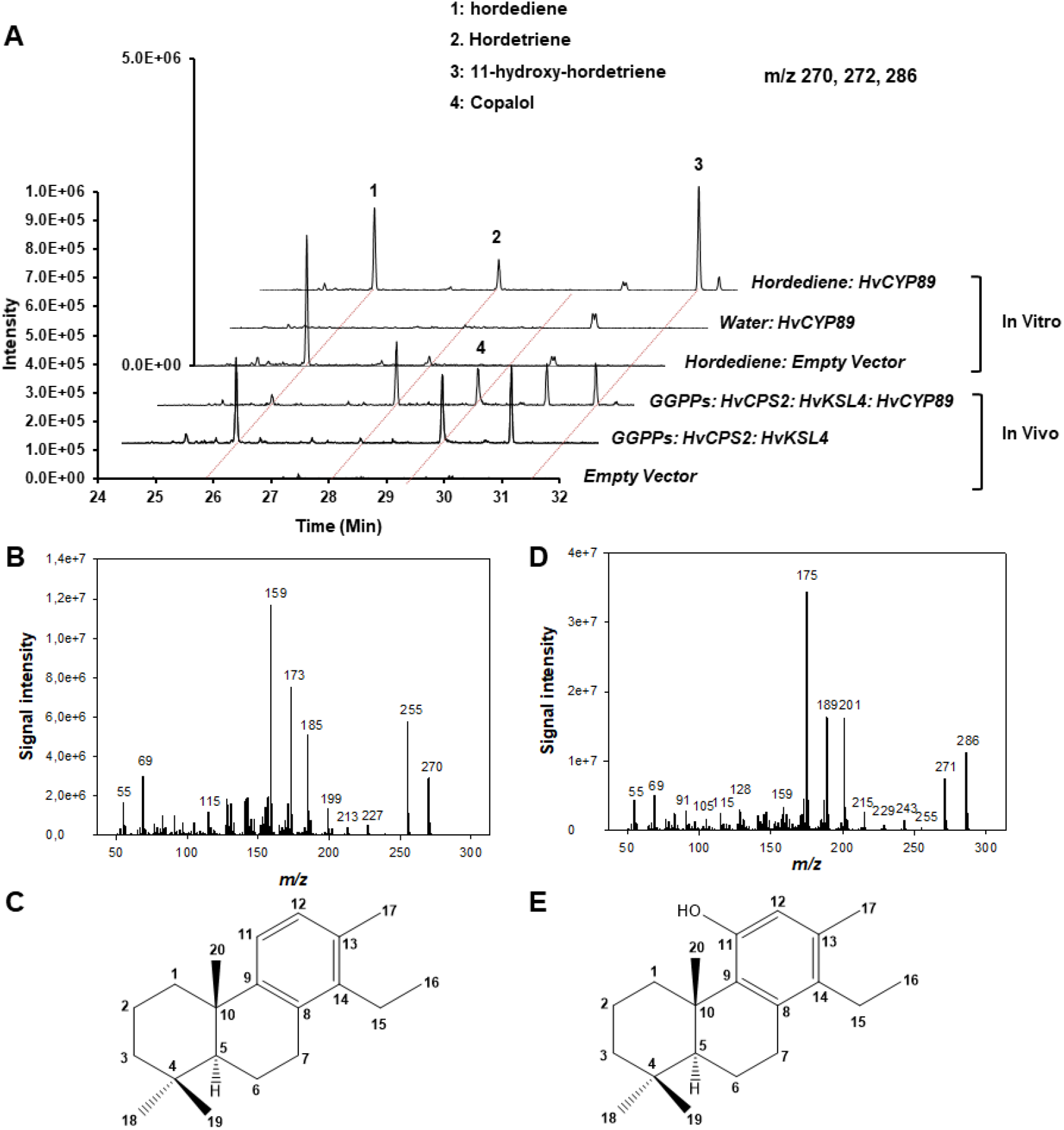
Characterization of HvCYP89E31. A. GC-MS chromatograms (single ion monitoring for *m/z* 270, 272 and 286). The in vivo chromatograms show analysis of extracts from transient assays in *N. benthamiana*. The in vitro chromatograms show analysis of in vitro assays with microsome fractions from yeast expressing HvCYP89E31. The identity of peaks is indicated by numbers. B and C. Respectively EI mass spectrum and structure of compound **2**, hordetriene. D and E. Respectively EI mass spectrum and structure of compound **3**, 11-hydroxy-hordetriene.

### Hordetriene and 11-hydroxy-hordetriene are present in barley roots infected with *B. sorokiniana*

To determine if the diterpene products identified in yeast and *N. benthamiana* by metabolic engineering are also produced in barley plants, we infected barley roots with the pathogen *B. sorokiniana*, or with the beneficial fungus *Serendipita vermifera* (syn. *Sebacina vermifera*) or with both as described in Sarkar et al. (2019). Hexane extracts from the roots were then analyzed by GC-MS and compared to an extract from a yeast strain expressing HvCPS2, HvKSL4 and CYP89E31 (**Fig. 5**). Whereas hordediene (**1**) could not be detected in any of the root extracts, **2** and **3** were detected in roots infected with *B. sorokiniana* or with *B. sorokiniana* and *S. vermifera*. In mock-infected roots or in roots infected with *S. vermifera* alone, smaller amounts of **3** could be detected compared to roots infected with *B. sorokiniana*, but **2** could not be detected. These results demonstrate that the diterpenoids produced by enzymes of the chromosome 2 cluster are indeed induced during fungal pathogen infection but not significantly by beneficial endophytic colonization, an observation consistent with the transcriptome data generated previously (Sarkar et al., 2019).

**Figure 5.**
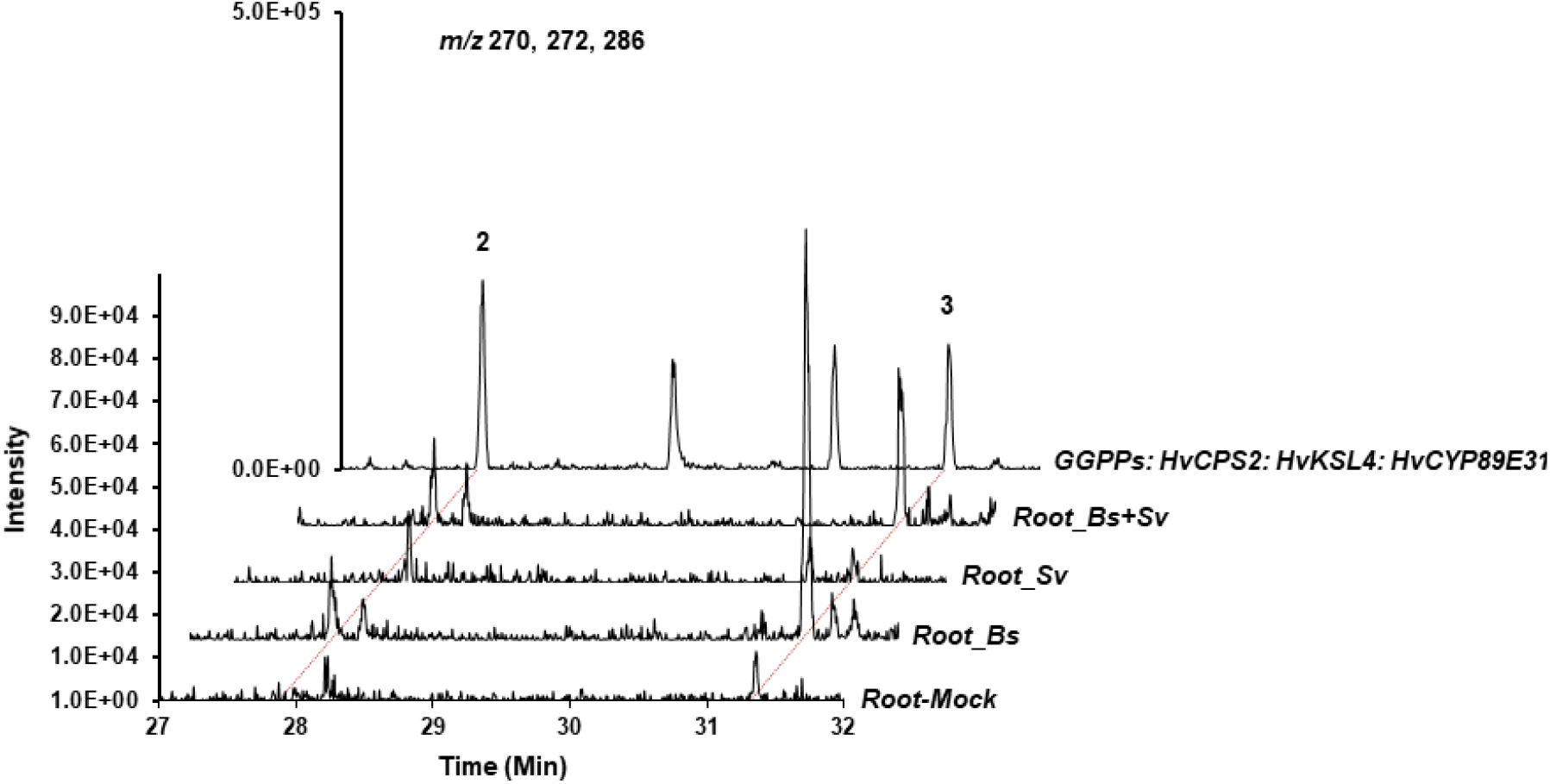
GC-MS chromatograms of extracts from barley roots infected with *B. sorokiniana* and/or *S. vermifera*. Single ion monitoring (m/z 270, 272 and 286) of extracts from roots infected with *B. sorokiniana* (Root_Bs), *S. vermifera* (Root_Sv) or both (Root_Bs+Sv) or mock inoculated (Root-Mock). An extract of yeast strain expressing HvCPS2, HvKSL4 and HvCYP89E31 is shown as a reference. **2**: hordetriene; **3:** 11-hydroxy-hordetriene.

### Detection/identification of diterpenoid phytoalexins induced by *B. sorokiniana* infection

Given the presence of additional CYP enzymes in the barley chromosome 2 cluster, we wondered whether further oxidized diterpenoids could be detected in barley roots infected with *B. sorokiniana*. We thus performed untargeted LC-QToF-MS (negative mode) analysis of extracts from the root and from the medium. Searching for masses corresponding to additional oxidations of **3 (Fig. 4)**, i.e. 301.2, 317.2, 315.2 and 331.2, a number of peaks could be detected with a strong increase in samples from roots and medium of plants infected with *B. sorokiniana* (**Fig. 6**). By comparison, under mock control conditions this accumulation was not observed. LC-TOF-MS with negative electrospray ionization detected [M-H] adducts of six major compounds of which two products with the monoisotopic neutral mass or 316.21 Da dominated. High enrichments were also observed for three compounds with the monoisotopic formula mass 332.20 Da and for one with 318.22 Da, indicating that these main products carry 3 or 4 oxygen atoms. MS/MS spectra of these compounds imply a diterpene character based on the fragment ions 269.19, 271.21, 287.20 and 301.1 (**Fig. S3**). Neutral losses of 43.99 and 46.01 suggest the presence of lactones or carboxyl groups in all six diterpenes. As well, compound 6 has an additional carbonyl group as indicated by a neutral loss of 30.01 Da between the molecular ion and m/z 287.203.

**Figure 6.**
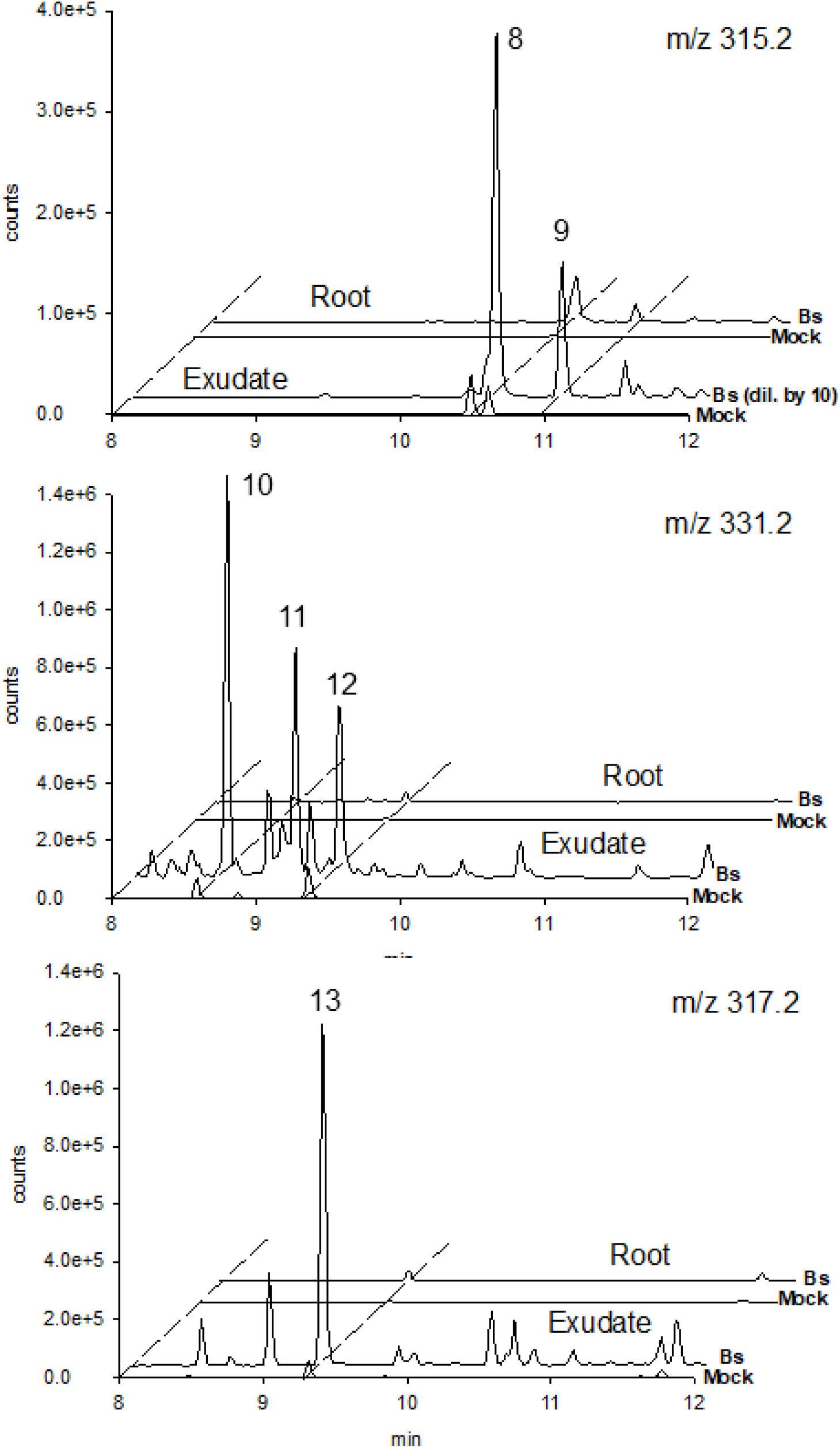
LC-(-)ESI-QToF-MS of major diterpene metabolites in roots and root exudates of barley 6 dpi with *B. sorokiniana* (Bs) and in respective mock infected controls. HR-MS/MS spectra of compounds 8-13 are presented in **Fig. S3**.

## Discussion

### Evolution and diversity of diterpenoid phytoalexins in monocots

Since the discovery of diterpenoid phytoalexins in rice and the elucidation of their biosynthesis, there has been an increasing number of reports on investigations of specialized diterpenoid metabolism in monocots. This includes maize (*Zea mays*) (Schmelz et al., 2011; Ding et al., 2020), switchgrass (*Panicum virgatum*) (Pelot et al., 2018; Muchlinski et al., 2021), and wheat (*Triticum aestivum*) (Wu et al., 2012; Zhou et al., 2012). In the Triticeae, which comprise wheat, barley and rye *(Secale cereale*), despite the publication of a report describing the characterization of wheat diterpene synthases, (Wu et al., 2012; Zhou et al., 2012), to the best of our knowledge no diterpenoid phytoalexins had been identified yet. Our work shows that like maize and rice, barley is also able to produce diterpenoid phytoaelxins when challenged with a pathogen. Thus, production of diterpenoid phytoalexins appears to be a common feature of monocots. It is also noteworthy that the genes involved in the biosynthesis of these compounds are frequently localized in clusters encoding diterpene synthases (CPS and KSL) as well as CYPs or in the case of rice a short chain dehydrogenase reductase (OsMAS) (Shimura et al., 2007; Ding et al., 2020; Muchlinski et al., 2021). Clusters of genes involved in specialized metabolism are encountered in plants and typically consist of tandem duplicated copies of one or more gene families (Nützmann et al., 2016). The molecular mechanisms driving the formation and evolution of these clusters are still enigmatic, but the physical association and duplication of genes offer a number of potential advantages. First, physical association means that possible toxic intermediates due to mutation or loss of one of the genes are less likely to accumulate. Second, the presence of multiple copies of highly similar genes provides opportunities for gene rearrangements and independent evolution, thereby opening the door to rapid chemical innovations. The capacity to evolve novel defenses rapidly constitutes a key evolutionary adaptation in the arms race against pests and pathogens.

Despite the relatedness of wheat and barley, the characterization of wheat diterpene synthases indicates that none of them can produce the cleistanthane backbone identified here (Awika, 2011). This is additional evidence that the gene clusters allow for rapid evolution of enzyme function and chemical diversification.

A search for regions in rice that are syntenic to the barley cluster on chromosome 2 using the SynFind tool (Tang et al., 2015) was unsuccessful (data not shown). This could be due to the fact that this chromosomal region is subject to frequent rearrangements. Nonetheless, in a publication investigating the synteny between rice and barley (Thiel et al., 2009), much of barley chromosome 2 was found to be syntenic with rice chromosome 4, which is where the main cluster for momilactone biosynthesis is located. Thus, it is possible that the evolution of this genomic cluster predates the divergence between Triticeae and rice.

### Hordediene: an unprecendented diterpene backbone in grasses

HvKSL4 catalyzes the cyclization of (+)-CPP to hordediene, a cyclization reaction which, to the best of our knowledge, has not been reported previously, at least not in monocots. Hordediene belongs to the cleistanthane group of labdane-related diterpenes, which is characterized by the presence of an ethyl group attached to C17, instead of C13 as in pimaradiene. In the cyclization pathway we propose, after the initial dephosphorylation, the first electron migrations lead to the formation of the pimaradienyl cation (**Fig. S4**). At this point the pathway diverges from the pimaradiene cyclization via capture of a proton from C9 by the C14-C15 double bond of pimaradienyl. This results in formation of the C8-C9 double bond and the C14 carbocation. A hydride shift from C17 to C14 would then allow the migration of the ethyl group to C17 and consequently result in the presence of carbocation at C13. Resolution of this cation by proton loss from C12 would then lead to hordediene.

A survey of the literature on cleistanthane diterpenoids reveals a modest number of reports (65 in Web of Science as of May 2021), with occurrence either in plants or fungi. Many of the cleistanthanes reported are of the *ent*- configuration, such as Fimbricalyxoid A from *Strophioblachia fimbricalyx* (Cheng et al., 2016) but some of normal (+) configuration, like hordediene, have also been described, such as zythiostromic acids (Ayer and Khan, 1996). There is little information on the biological function of these diterpenoids, although in some cases anticancer activity in the μM range could be determined (Cheng et al., 2016).

### Production and potential role of diterpenoid phytoalexins in barley

We characterized three genes from the cluster (HvCPS2, HvKSL4 and HvCYP89E31) and showed that co-expression of these three genes results in the production of two major products, hordetriene and 11-hydroxy-hordetriene. The fact that we could detect both of these products in barley roots infected with *B. sorokiniana*, strongly supports a role of this cluster in the production of diterpenoid phytoalexins and in the defense reaction of barley. LC-MS analysis indicated the presence of further oxidized diterpenoids, however at this stage we were not able to isolate and purify sufficient quantity of these compounds for structural elucidation. Thus it remains to be determined whether these are also derived from the same diterpene backbone as compounds **2** and **3**. The fact that no other diterpene synthase is induced by *B. sorokinianasorokiniana* in our transcriptome data argues in favor of this hypothesis. At this stage, we can only speculate on the biological activity of these compounds. Their biosynthesis is only minimally induced by *S. vermifera*, but very strongly by the pathogenic fungus *B. sorokiniana* (Sarkar et al., 2019). Whether the induction is specific to *B. sorokiniana* or is a general response to infection by pathogens remains to be determined. Interestingly, in a recent report on the metabolic profiling of barley plants infected with *Fusarium graminearum*, which causes head blight, two diterpenoids with masses of 302.2 and 318.2 were the most strongly induced compounds (respectively 32 and 22 fold change) (Karre et al., 2017). They were annotated as neoabietic and 7-hydroxykaurenoic acid respectively, although there was no formal identification of these compounds. We suspect that these compounds also belong to the diterpenoid group that we identified, which would suggest that they are not induced specifically by *B. sorokiniana*. Availability of larger amounts of pure compounds in the future will allow us to perform bioassays and determine their potential antimicrobial activity. Of relevance to their biological function is the fact that these compounds are for the most part secreted into the medium by the roots. Since they are present at a low basal level in non-infected plants, it is likely that their presence in the rhizosphere could influence the composition of the root microbiome. There is already evidence of specialized metabolites secreted by the roots that impact the composition of the microbiome (Massalha et al., 2017; Huang et al., 2019; Murphy et al., 2021). Conversely, specific members of the microbial community can induce secretion of specialized metabolites by the roots (Massalha et al., 2017), suggesting a complex network of signals and effectors between the microbial community and the plant in the rhizosphere. Generating barley plants mutated in HvCPS2 or HvKSL4 by CRISPR-Cas gene editing will allow us to address these questions in this important crop species in the future.

## Materials and Methods

### Plant growth and fungal inoculations

Barley seeds (*Hordeum vulgare* L. cv Golden Promise) were sterilized in 70% ethanol for 1 min, followed by washing with sterile distilled water and 1.5 h incubation in 12% sodium hypochloride under continuous shaking. After 3 times 30 min washing, the seeds were placed on wet filter paper in darkness and at room temperature for 4 days for germination. Four seedlings were transferred to 1/10 PNM (Plant Nutrition Medium, pH 5.7) (Wawra et al., 2016) in sterile glass jars and grown in a day/night cycle of 16/8 h at 22/18 °C, 60 % humidity under 108 μmol m^−2^ s^−1^ light intensity.

*Bipolaris sorokiniana* (ND90Pr) and *Serendipita vermifera* (MAFF305830) were used in this study. *Bs* was propagated on modified CM medium with 1.5% agar and *Sv* on MYP medium with 1.5% agar in the dark at 28 °C for 21 days and 14 days before inoculation respectively. *Bs* conidia and *Sv* mycelial were collected according to the procedures which were described in (Sarkar et al., 2019). Barley roots were inoculated with 3 ml of either *Sv* mycelium (2g/50ml), *Bs* conidia (5000 spores/ml) or a 1:1 mixture of the two fungi per jar for 6 days. Sterile water was used as a mock treatment. Roots washed thoroughly and the corresponding medium were collected and snap-frozen in liquid nitrogen for extraction of metabolites.

### qRT-PCR

RNA isolation from roots was performed using the Spectrum Plant Total RNA kit (Sigma-Aldrich). The complementary DNA (cDNA) was synthesized using ProtoScript II First Strand cDNA Synthesis Kit (New England Biolabs) following the manufacturer’s instructions with primer d(T)_23_ VN. Quantitative real-time PCR was performed in triplicates using 10-20 ng cDNA as template and gene specific primer pairs shown in **Table S8** in CFX Connect Real-Time PCR System (Bio-Rad). The PCR conditions were 95◻°C for 15◻min; 40 cycles of 95 °C for 15 s, 56◻°C for 30◻s; 95 °C for 10 s. The melting curve was measured from 65 °C to 95 °C with a step of 0.1◻°C per second. Relative expression of targeted genes was calculated using delta Ct method (Livak and Schmittgen, 2001) and barley ubiquitin genes as references (Deshmukh et al., 2006).

### Phylogenetic analysis

Amino acid sequences (**Tables S2, S3 and S5**) were aligned and phylogenetic trees were generated using MEGA X (Kumar et al., 2018). ClustalW was used for the alignment and the maximum likelihood method with a bootstrap of 1,000 for the phylogenetic tree. For the other parameters, default settings were used.

### Heterologous expression of diterpenes in yeast

Plasmids containing *GGPPS* and *ATR1* were kindly provided by colleagues in the group and have been described previously (Scheler et al., 2016). Codon-optimized DNA sequences of *HvCPS2, HvKSL4* and *HvCYP89E31* were synthesized by Thermo Fisher Scientific Inc. for yeast expression. Each gene was further cloned into Golden Gate compatible yeast expression level 1 vector, together with a synthetic galactose-inducible promoter and a terminator. Different gene combinations were finally assembled into one yeast expression level M vector by a 50 cycle restriction–ligation reaction with BpiI and T4-Ligase. Constructs were then transformed into *S. cerevisiae* strain INVSc1 (Thermo Fisher Scientific Inc.) and plated out onto uracil-free (Ura-) selection medium (1◻g◻l^−1^ Yeast Synthetic Drop-out Medium Supplements without uracil (Sigma-Aldrich), 6.7◻g◻l^−1^ Yeast Nitrogen Base with Amino Acids (Sigma-Aldrich) and 20◻g◻l^−1^ Micro Agar (Duchefa Biochemie)). Three positive colonies were picked and inoculated into 5◻ml yeast extract-peptone-dextrose (YPD) medium (20◻g◻l^−1^ tryptone and 10◻g◻l^−1^ yeast extract) containing 2% of glucose and grown for 24◻h with shaking at 30◻°C. To induce protein expression, the cell pellet was resuspended in fresh YPD medium containing 2% galactose. After another 24◻h of growth, the whole culture was extracted with 2◻ml *n*-hexane.

### Transient expression in *Nicotiana benthamiana*

Transit peptides of protein HvCPS2 and HvKSL4 were predicted by two online tools, ChloroP 1.1 (http://www.cbs.dtu.dk/services/ChloroP/) and LOCALIZER (http://localizer.csiro.au/). Truncated sequences without the predicted transit peptides of these two genes were generated by PCR reactions using designed primers and subsequently sequenced. The cDNAs of *HMG reductase*, *GGPPS* in plasmids have been described previously (Scheler et al., 2016; Yadav et al., 2019). The *HMG reductase*, *GGPPS*, *trHvCPS2*, *trHvKSL4* and *HvCYP89E31* were cloned into T-DNA vectors (binary vector pL1F-1) driven by the 35S promoter and flanked by the Ocs terminator (Weber et al., 2011). The resulting T-DNA plasmids were transformed into *Agrobacterium tumefaciens* strain GV3101::pMP90 and plated out onto LB agar plates with appropriate antibiotics. Bacteria were harvested and resuspended in infiltration medium (10 mM MgCl_2_, 10 mM MES, 20 μM acetosyringone, pH=5.6) after 48 h inoculation at 28◻°C. To co-infiltrate several genes, each bacteria suspension was diluted to a final OD_600_ of 0.4, then all strains were mixed equally to an appropriate volume for infiltration. The suspension was infiltrated into the abaxial side of several leaves in three individual 4-week-old *N. benthamiana* plants using a syringe without needle. After treatment, the plants were cultivated in a climate controlled phytochamber for 4 days. Three leaf discs (9 mm diameter) per infiltrated spot were harvested and extracted by 2 ml *n*-hexane, followed by drying down under nitrogen flow and GC-MS analysis.

### Microsome isolation and *in vitro* enzyme assay

A protocol from the literature with slight modification was used for microsome isolation (Urban et al., 1997; Scheler et al., 2016). The construct carrying *HvCYP89E31* and *ATR1* were transformed into yeast strain INVSc1. A single positive colony was picked to inoculate 5◻ml of Ura-medium with 2% glucose and grown for 24◻h at 30◻°C with shaking. The culture was then used to inoculate 100◻ml of Ura-medium with 2 % glucose in a 500◻ml flask at 30◻°C for 24◻h. The cells were then collected by centrifugation, resuspended in 100◻ml fresh YPD medium with 2 % galactose to induce protein expression and inoculated under shaking for another 24◻h at 30◻°C. All the following steps were carried out at 4◻°C. The cells were harvested by centrifugation and resuspended in 30 ml of pre-chilled TEK buffer (50◻mM Tris-HCl pH 7.5, 1◻mM EDTA, 100◻mM KCl), centrifuged again and resuspended in 2◻ml TES buffer (50◻mM Tris-HCl pH 7.5, 600◻mM sorbitol, 10◻g◻l^−1^ BSA, 1.5◻mM β-mercaptoethanol) and transferred to a 50◻ml tube. Acid-washed autoclaved 450– 600◻μm diameter glass beads were added into the tube until the surface of the cell suspension are reached. The suspension was shaken vigorously by hand for 1◻min and returned back to ice for 1◻min. This step was repeated four times. The glass beads were washed by 5◻ml TES buffer three times, and the supernatant was collected and combined to a new tube, followed by centrifugation at 7,500◻g for 10◻min. The supernatant was transferred to ultracentrifugation tubes and centrifuged for 2◻h at 100◻000◻g. The pellet was gently washed successively with 5◻ml TES and 2.5◻ml TEG buffer (50◻mM Tris-HCl pH 7.5, 1◻mM EDTA and 30% glycerol) after the supernatant was removed, then transferred to a Potter homogenizer with a spatula. 2 ml TEG buffer was added to the homogenizer and the pellet was carefully homogenized. 100 μl aliquots were transferred to 1.5◻ml microtubes and stored at −80◻°C until used.

In vitro CYP enzyme assays were performed in a 600◻μl reaction volume, containing 40◻μl of microsome preparation, 100◻μM substrate, 1◻mM NADPH, 50◻mM sodium phosphate pH 7.4. The solution was incubated at 30◻°C for 2◻h with gentle shaking. Products were extracted with 1◻ml *n*-hexane under strong agitation (vortex). After centrifugation, the organic phase was collected, then dried under a N_2_ stream and resuspended in 100◻μl *n*-hexane for GC-MS analysis.

### Purification of diterpenes by silica gel column chromatography or SPE

For the purification of hordediene, 1 l of yeast culture was grown and extracted with 1 l *n*-hexane. The raw extracts were dried in a rotary evaporator and resuspended in 4 ml n-hexane, then loaded into two properly conditioned SiOH SPE cartridges (500 mg, MACHEREY-NAGEL). The cartridges were then washed with 2 ml n-hexane. The breakthrough and washing fraction were collected and combined. After drying down under nitrogen stream, an aliquot was measured by GC-MS to check the purity of the product and the rest, with an amount of around 2 mg was used for NMR structure elucidation.

Hordetriene and 11-hydroxy-hordetriene were first extracted from three liters of yeast culture. After concentration, the raw extracts were dissolved in 4 ml *n*-hexane and loaded into a pre-conditioned self-packed silica gel column (5 g, 15 mm × 100 mm). The column was then eluted by *n*-hexane and successively by 99:1, 98:2, 97:3, 96:4, 96:5 n-hexane: ethyl acetate solutions. The volume of each elution solution was 10 ml but the elution was separately collected in five 2 ml microtubes. An aliquot of each fraction was measured by GC-MS and the fractions with the same product were combined and then used for NMR structure elucidation. The yield of hordetriene and 11-hydroxyhordetriene was around 0.5 mg.

### Nuclear magnetic resonance (NMR) conditions

H, ^13^C, 2D (^1^H,^1^H gDQCOSY; ^1^H,^1^H zTOCSY;^1^H,^1^H ROESYAD; ^1^H,^13^C gHSQCAD; 1H,^13^C gHMBCAD), selective (^1^H,^1^H zTOCSY1D; ^1^H,^1^H ROESY1D), and band selective (^1^H,^13^C bsHMBC) NMR spectra were measured with an Agilent VNMRS 600 instrument at 599.83 MHz (^1^H) and 150.84 MHz (^13^C) using standard CHEMPACK 8.1 pulse sequences implemented in the VNMRJ 4.2A spectrometer software. TOCSY mixing time: 80 ms; ROESY mixing time: 300 ms; HSQC optimized for ^1^J_CH_ = 146 Hz; HMBC optimized for ^n^J_CH_ = 8 Hz. All spectra were obtained with C_6_D_6_ + 0.03% TMS as solvent at +25°C. Chemical shifts were referenced to internal TMS (δ ^1^H = 0 ppm) and internal C_6_D_6_ (δ ^13^C = 128.0 ppm).

### Metabolites extraction from barley roots and PNM medium

100 mg (fresh weight) of frozen and cryo-ground root matter was extracted using 900 μL dichloromethane/ethanol (2:1, v/v) and 100 μl hydrochloric acid solution (pH 1.4). Extraction and duplicate removal of hydrophilic metabolites was achieved by 1 min FastPrep bead milling (FastPrep24, MP Biomedicals) followed by phase separation during centrifugation. For extraction 1.6 mL wall-reinforced cryo-tubes (Biozyme) each containing steel and glass beads were used. The upper aqueous phase was discarded and replaced for a second round of bead mill extraction/ centrifugation. Thereafter, the aqueous phase was removed and the lower organic phase was collected. Subsequently 600 uL tetrahydrofuran (THF) was used for exhaustive extraction (FastPrep). After centrifugation the organic THF extract was combined with the first extract and dried under a stream of N_2_.

Root exudates were extracted from 60 mL of gelrite media. For this, the gel was distributed into two 50 ml Falcon tubes. To each tube 4 g of NaCl and 3 mL ethyl acetate were added. The tubes were thoroughly shaken by hand and centrifuged. The upper phase (organic extract) was collected before fresh ethyl acetate was added for another two consecutive extractions. The combined extracts of three extraction rounds were combined and dried in a stream of N_2_.

### GC-MS

Dried extracts were suspended in 200 μl n-hexane. The analysis of yeast and plant extracts was carried out using a Trace GC Ultra gas chromatograph (Thermo Scientific) coupled to an ATAS Optic 3 injector and an ISQ single quadrupole mass spectrometer (Thermo Scientific) with electron impact ionization. Chromatographic separation was performed on a ZB-5ms capillary column (30◻m × 0.32◻mm, Phenomenex) using splitless injection and an injection volume of 1◻μl. The injection temperature rose from 60◻°C to 250◻°C with 10◻°C◻s^−1^ and the flow rate of helium was 2◻ml◻min^−1^. The GC oven temperature ramp was as follows: 50◻°C for 1◻min, 50 to 300◻°C with 7◻°C◻min^−1^, 300–330◻°C with 20◻°C◻min^−1^ and 330◻°C for 5◻min. Mass spectrometry was performed at 70◻eV, in a full scan mode with *m/z* from 50 to 450. Data analysis was done with the device specific software Xcalibur (Thermo Scientific).

### RP-UPLC-ESI-MS/MS

For UPLC-MS analysis dried extracts were suspended in 100 uL 80% methanol/ 20% water. Separation of medium polar metabolites was performed on a Nucleoshell RP18 (2.1 × 150 mm, particle size 2.1 μm, Macherey & Nagel, GmbH, Düren, Germany) using a Waters ACQUITY UPLC System, equipped with a Binary Solvent Manager and Sample Manager (20 μl sample loop, partial loop injection mode, 5 μl injection volume, Waters GmbH Eschborn, Germany). Eluents A and B were aqueous 0.3 mmol/L NH_4_HCOO (adjusted to pH 3.5 with formic acid) and acetonitrile, respectively. Elution was performed isocratically for 2 min at 5% eluent B, from 2 to 19 min with a linear gradient to 95% B, from 19-21 min isocratically at 95% B, and from 21.01 min to 24 min at 5% B. The flow rate was set to 400 μl min^−1^ and the column temperature was maintained at 40 °C. Metabolites were detected by positive and negative electrospray ionization and mass spectrometry.

Mass spectrometric analysis of small molecules was performed by MS-TOF-SWATH-MS/MS (TripleToF 5600, AB Sciex GmbH, Darmstadt, Germany) operating in negative or positive ion mode and controlled by Analyst 1.7.1 software (AB Sciex GmbH, Darmstadt, Germany). The source operation parameters were as follows: ion spray voltage, −4500 V / +5500 V; nebulizing gas, 60 psi; source temperature 600°C; drying gas, 70 psi; curtain gas, 35 psi. TripleToF instrument tuning and internal mass calibration were performed every 5 samples with the calibrant delivery system applying APCI negative or positive tuning solution, respectively (AB Sciex GmbH, Darmstadt, Germany).

TripleToF data acquisition was performed in MS1-ToF mode and MS2-SWATH mode. For MS1 measurements, ToF masses were scanned between 65 and 1250 Dalton with an accumulation time of 50 ms and a collision energy of 10V (−10V). MS2-SWATH-experiments were divided into 26 Dalton segments of 20 ms accumulation time. Together the SWATH experiments covered the entire mass range from 65 to 1250 Dalton in 48 separate scan experiments, which allowed a cycle time of 1.1 s. Throughout all MS/MS scans a declustering potential of 35 (or −35 V) was applied. Collision energies for all SWATH-MS/MS were set to 35 V (−35) and a collision energy spread of ±25V, maximum sensitivity scanning, and elsewise default settings.

## Acknowledgments

This work was funded by grant TI 800/7-1 from the *Deutsche Forschungsgemeinschaft* (DFG) to AT and by grant ZU 263/11-1 to AZ. This project is part of the Priority Programme of the DFG SPP2125 “Deconstruction and Reconstruction of the Plant Microbiota, DECRyPT).” (https://ag-zuccaro.botanik.uni-koeln.de/decrypt).

## Author Contributions

YL performed the LC-MS, GC-MS, yeast and *N. benthamiana* expression and gene expression analysis. GB provided supervision for LC-MS and data analysis. AS-H provided assistance for LC-MS sample preparation. AP performed the NMR measurements and analyses. UB supervised the yeast expression experiments. LM and AZ provided infected barley samples and transcriptomic data. AT and GB designed the project. AT designed and supervised the project. AT wrote the abstract, introduction, results and discussion. YL wrote the materials and methods, except the NMR part (AP) and LC-MS (GB). AT, YL and GB prepared the figures. All authors read and approved the manuscript.

