## Supplemental Information for "A barley gene cluster for the biosynthesis of diterpenoid phytoalexins"

**Figure S1. GC-MS analysis of transient expression in *N. benthamiana*.**

**Figure S2. Phylogenetic analysis of CYP sequences from barley chromosome 2 diterpenoid cluster.**

**Figure S3. High resolution MS/MS spectra of 6 diterpenoids produced by Barley Cv. Golden Promise after infection with the pathogen *B. sorokiniana.***

**Figure S4. Proposed cyclization pathway of (+)-copalyl diphosphate to hordediene by HvKSL4.**

**Table S1. World production and area harvested for barley in 2019.**

**Table S2. CPS sequences used for phylogenetic analysis in Figure 2.**

**Table S3. KS and KSL sequences used for phylogenetic analysis in Figure 2.**

**Table S4. NMR data of hordediene (compound 1)**

**Table S5. List of sequences used for the phylogenetic analysis of CYP sequences.**

**Table S6. NMR data of hordetriene (compound 2)**

**Table S7. NMR data of 11-hydroxy-hordetriene (compound 3)**

**Table S8. Oligonucleotides used in this study.**

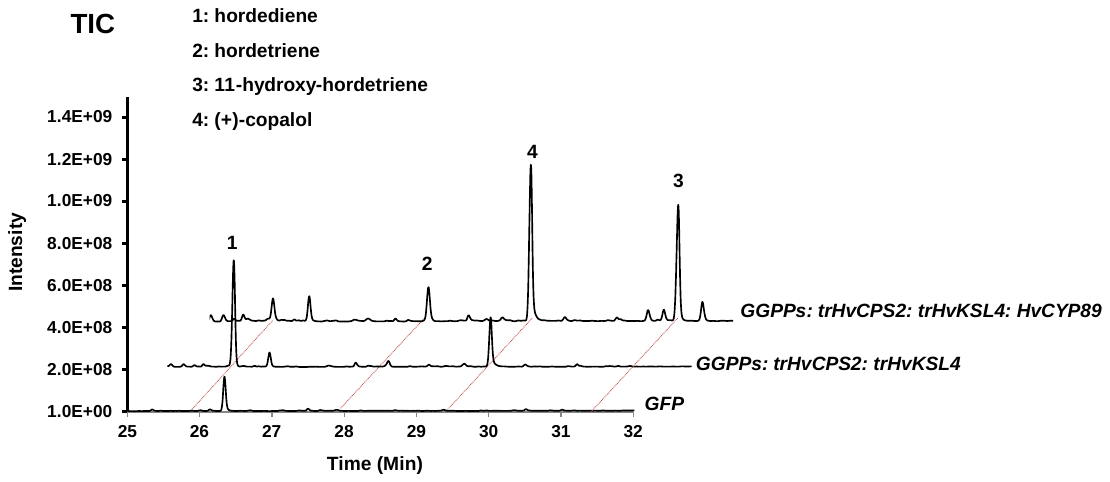

**Figure S1. GC-MS analysis of transient expression in *N. benthamiana*.** The genes that were infiltrated are indicated ion the right side next to the corresponding chromatogram. Shown are total ion chromatograms (TIC). Substances detected are indicated by numbers.

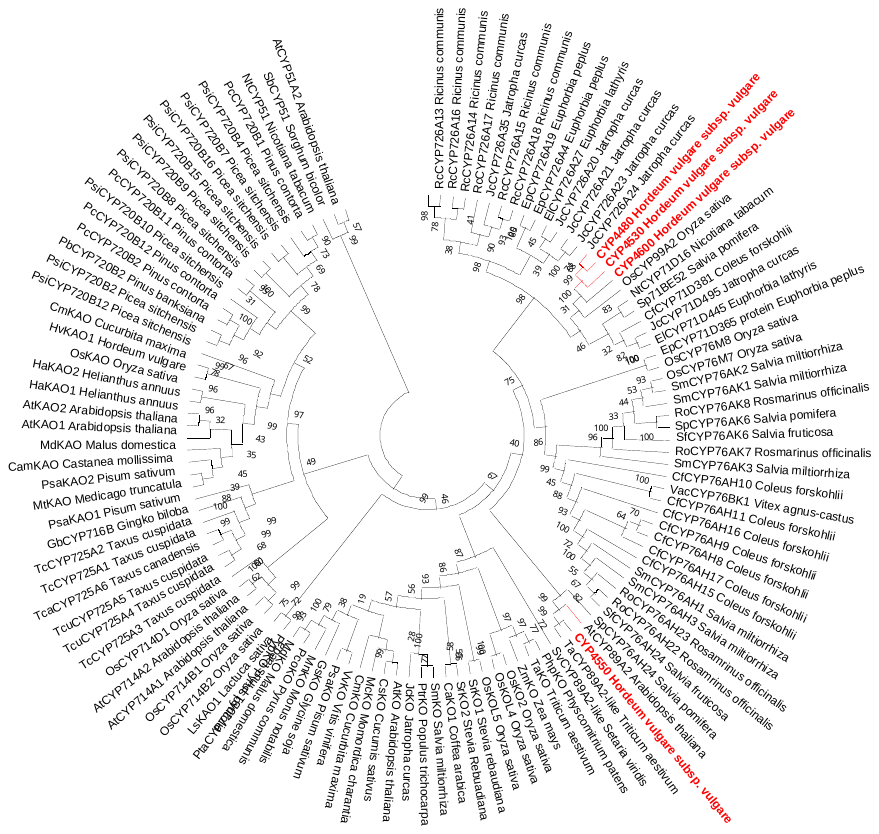
**Figure S2. Phylogenetic analysis of CYP sequences from barley chromosome 2 diterpenoid cluster.** The evolutionary history was inferred by using the Maximum Likelihood method and Poisson correction model [1]. The bootstrap consensus tree inferred from 1000 replicates [3] is taken to represent the evolutionary history of the taxa analyzed [3]. Branches corresponding to partitions reproduced in less than 50% bootstrap replicates are collapsed. The percentage of replicate trees in which the associated taxa clustered together in the bootstrap test (1000 replicates) are shown next to the branches [3]. Initial tree(s) for the heuristic search were obtained automatically by applying Neighbor-Join and BioNJ algorithms to a matrix of pairwise distances estimated using a JTT model, and then selecting the topology with superior log likelihood value. A discrete Gamma distribution was used to model evolutionary rate differences among sites (4 categories (+*G*, parameter = 2.6585)). This analysis involved 117 amino acid sequences. All positions with less than 95% site coverage were eliminated, i.e., fewer than 5% alignment gaps, missing data, and ambiguous bases were allowed at any position (partial deletion option). There were a total of 382 positions in the final dataset. Evolutionary analyses were conducted in MEGA X [2]. The list of sequences used is provided in **Table S5**. The CYPs from barley are in red.

1. Zuckerkandl E. and Pauling L. (**1965**). Evolutionary divergence and convergence in proteins. Edited in *Evolving Genes and Proteins* by V. Bryson and H.J. Vogel, pp. 97-166. Academic Press, New York.

2. Kumar S., Stecher G., Li M., Knyaz C., and Tamura K. (**2018**). MEGA X: Molecular Evolutionary Genetics Analysis across computing platforms. *Molecular Biology and Evolution* **35**:1547-1549.

3. Felsenstein J. (**1985**). Confidence limits on phylogenies: An approach using the bootstrap. *Evolution* **39**:783-791.

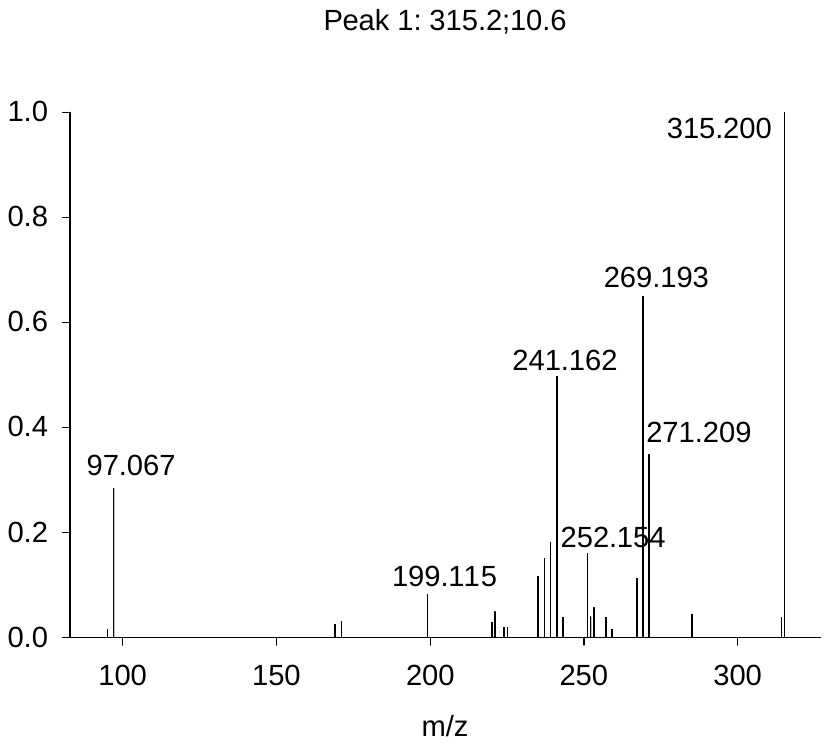

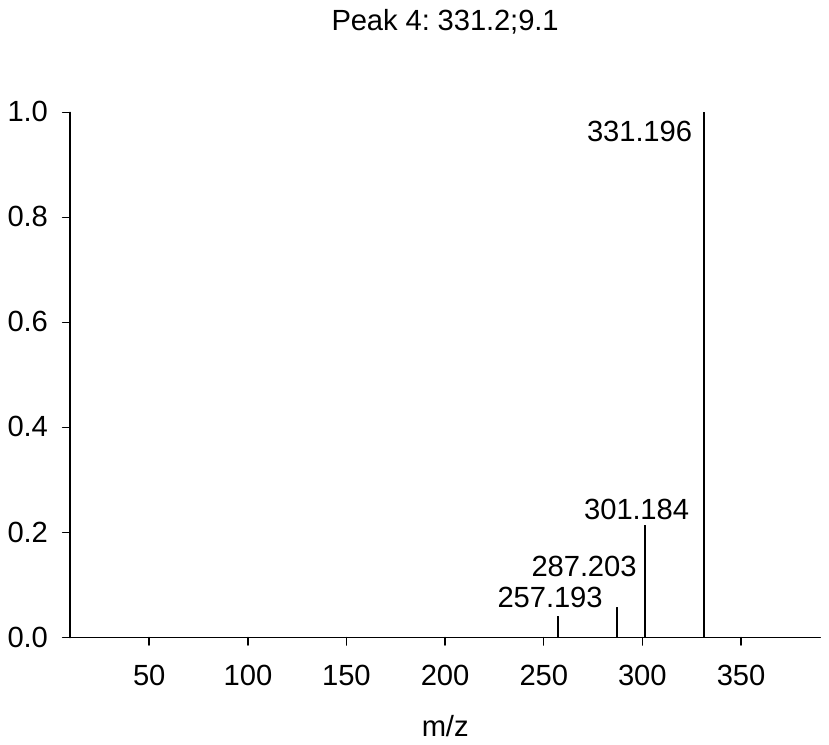

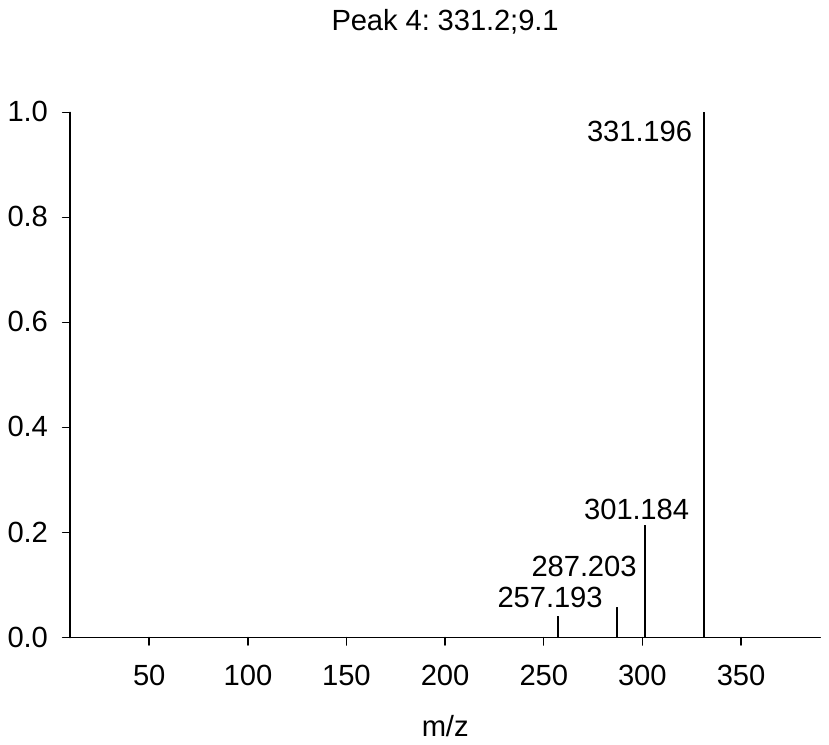

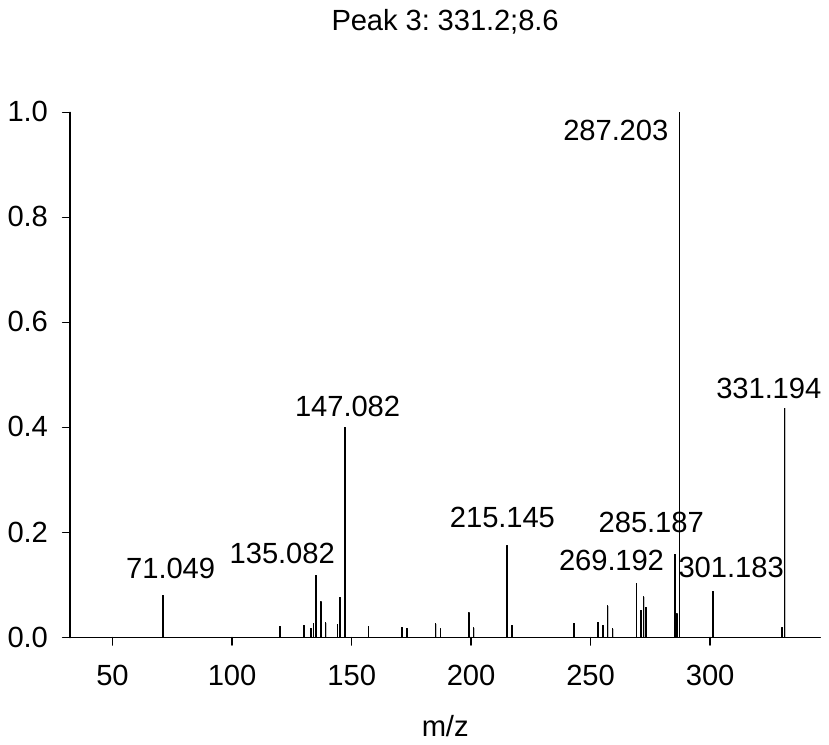

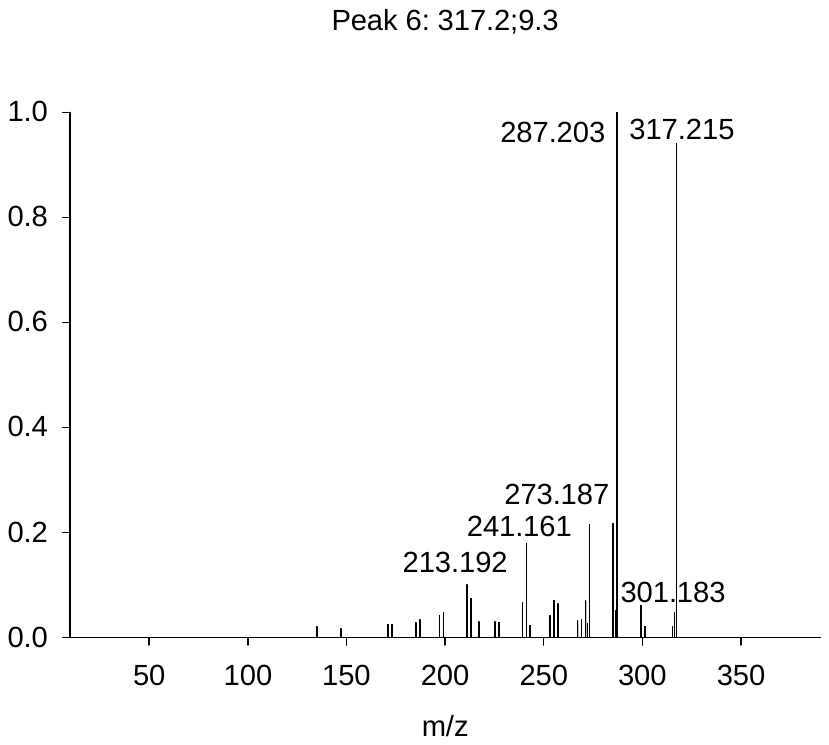

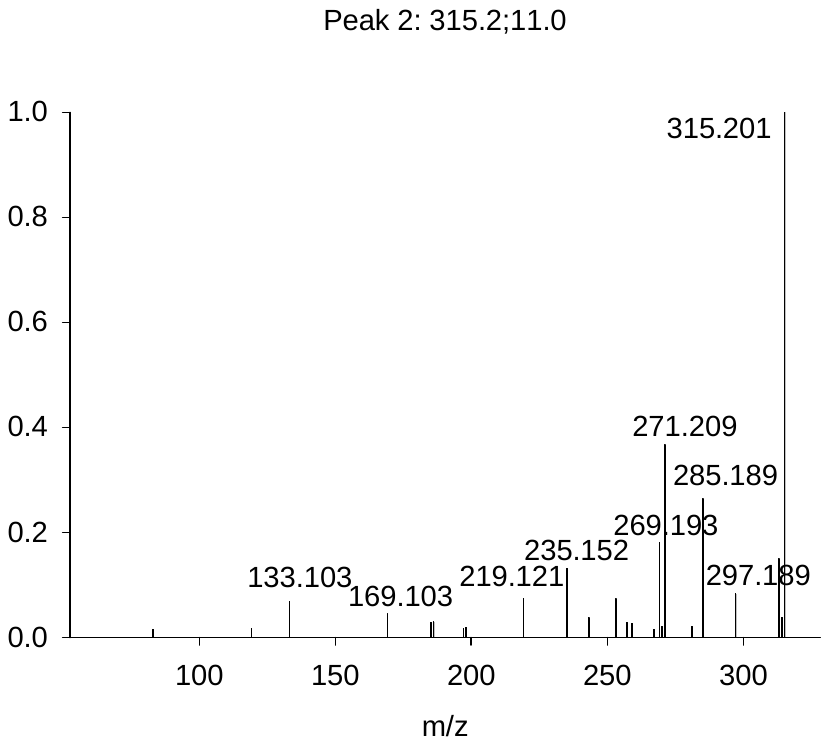

Compound 8: 315.2;10.6

Compound 9: 315.2;11.0

Compound 10: 331.2;8.6

Compound 11: 331.2;9.1

Compound 12: 331.2;9.4

Compound 13: 317.2;9.3

**Figure S3. High resolution MS/MS spectra of 6 diterpenoids produced by Barley Cv. Golden Promise after infection with the pathogen *B. sorokiniana.***

**Figure S4. Proposed cyclization pathway of (+)-copalyl diphosphate to hordediene by HvKSL4.**

**Table S1. World production and area harvested for barley in 2019. Source: FAO statistics.**

| **Area** | **Element** | **Item** | **Year** | **Unit** | **Value** | **Flag Description** |
| --- | --- | --- | --- | --- | --- | --- |
| World | Area harvested | Barley | 2019 | ha | 51 149 869 | Aggregate, may include official, semi-official, estimated or calculated data |
| World | Production | Barley | 2019 | tonnes | 158 979 610 | Aggregate, may include official, semi-official, estimated or calculated data |

**Table S2. CPS sequences used for phylogenetic analysis in Figure 2.**

| **Protein name** | **Species** | **Protein ID** |
| --- | --- | --- |
| AtCPS | *Arabidopsis thaliana* | NP_192187 |
| CmCPS1 | *Cucurbita maxima* | AAD04292 |
| OsCPS1 | *Orzya sativa* | BAF08464 |
| OsCPS2 | *Orzya sativa* | BAH91759 |
| OsCPS4 | *Orzya sativa* | NP_001052171 |
| TaCPS1 | *Triticum aestivum* | BAH56558 |
| TaCPS2 | *Triticum aestivum* | BAH56559 |
| TaCPS3 | *Triticum aestivum* | BAH56560 |
| TaCPS4 | *Triticum aestivum* | BAP01383 |
| HvCPS1 | *Hordeum vulgare* | AAT49065 |
| ZmCPS1 | *Zea mays* | NP_001105329 |
| ZmCPS2 | *Zea mays* | NP_001105257 |
| ZmCPS3 | *Zea mays* | AFW57228 |
| ZmCPS4 | *Zea mays* | AFW60403 |
| HvCPS2 | *Hordeum vulgare* | BAJ95441 |

**Table S3. KS and KSL sequences used for phylogenetic analysis in Figure 2.**

| **Protein name** | **Species** | **Protein ID** |
| --- | --- | --- |
| AtKS | *Arabidopsis thaliana* | AAC39443 |
| CmKS | *Cucurbita maxima* | Q39548 |
| OsKS | *Orzya sativa* | NP_001053841 |
| OsKSL4 | *Orzya sativa* | NP_001052175 |
| OsKSL5 | *Orzya sativa* | NP_001047190 |
| OsKSL6 | *Orzya sativa* | ABH10733 |
| OsKSL7 | *Orzya sativa* | NP_001047186 |
| OsKSL8 | *Orzya sativa* | NP_001067887 |
| OsKSL10 | *Orzya sativa* | NP_001066799 |
| OsKSL11 | *Orzya sativa* | Q1AHB2 |
| TaKS | *Triticum aestivum* | BAL41693 |
| TaKSL1 | *Triticum aestivum* | BAL41688 |
| TaKSL2 | *Triticum aestivum* | BAL41689 |
| TaKSL3 | *Triticum aestivum* | BAL41690 |
| TaKSL4 | *Triticum aestivum* | BAL41691 |
| HvKS | *Hordeum vulgare* | AAT49066 |
| ZmKSL1 | *Zea mays* | AFW61735 |
| ZmKSL2 | *Zea mays* | DAA54948 |
| ZmKSL3 | *Zea mays* | DAA36069 |
| ZmKSL4 | *Zea mays* | DAA49845 |
| HvKSL4 | *Hordeum vulgare* | BAK01991 |

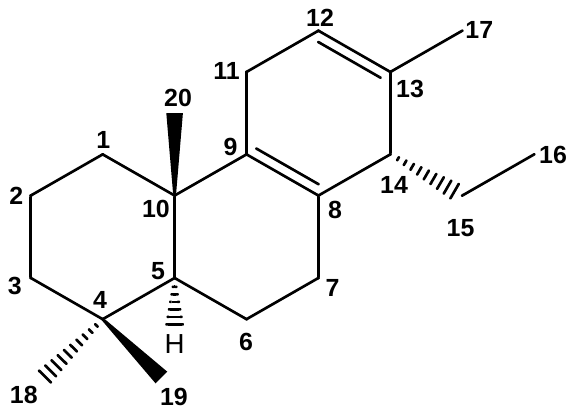
**Table S4. NMR data of hordediene (compound 1) (C_6_D_6_)**

| **Pos.** | **^13^C: δ [ppm]** | **^1^H: δ [ppm] m (J[Hz])** | **selected  HMBC (H 🡪 C)** | **ROE^c^** |
| --- | --- | --- | --- | --- |
| 1 | 37.38 | 1.66^a^ β; 1.043 ddd (13.0/13.0/3.7) α |  | 20; |
| 2 | 19.37 | 1.565^b^ α; 1.40^a^ β |  |  |
| 3 | 42.01 | 1.39^a^ β; 1.148 ddd (13.7/13.2/4.1) α |  | 18^d^ |
| 4 | 33.39 | --- |  |  |
| 5 | 52.49 | 1.210 dd (12.5/2.0) α |  | 1α^d^,7α^d^,18^d^ |
| 6 | 19.20 | 1.687^b^ α; 1.503 m β | 10; 5,7,10 | 18; |
| 7 | 30.81 | 2.189 m α; 1.750 br dd (17.1/6.2) β |  | 5^d^,15^d^,16^d^,18^d^; 14^d^ |
| 8 | 126.41 | --- |  |  |
| 9 | 138.22 | --- |  |  |
| 10 | 37.86 | --- |  |  |
| 11 | 26.41 | 2.53^a^ |  | 20 |
| 12 | 121.84 | 5.611 m |  | 11^d^,17^d^ |
| 13 | 133.66 | --- |  |  |
| 14 | 46.64 | 2.462 m | 8,9,12,13,15 | 7α^d^,7β^d^,15^d^ |
| 15 | 21.41 | 1.61^a^ |  |  |
| 16 | 22.89 | 0.763 t (7.4) | 14,15 |  |
| 17 | 7.72 | 1.656 qd (1.7/0.5) | 12,13,14 |  |
| 18 | 33.44 | 0.921 s | 3,4,5,19 |  |
| 19 | 21.94 | 0.877 s | 3,4,5,18 | 20 |
| 20 | 19.42 | 1.014 s | 1,5,9,10 |  |

^a^ ^1^H chemical shift of HSQC correlation peaks; ^b^ ^1^H chemical shift from selective TOCSY spectrum; ^c^ each NOE is listed only once; ^d^ correlations (H 🡪 H) from selective ROESY spectra. 14*S* configuration is more likely because of NOE H-7β/H-14 und H-7α/H-15.

**Table S5. List of sequences used for the phylogenetic analysis of CYP sequences.**

| **Accession no.** | **Name** | **Organism** | **Function** |
| --- | --- | --- | --- |
| AB014459 | AtCYP51A2 | *Arabidopsis thaliana* |  |
| Q93Z79 | AtCYP714A1 | *Arabidopsis thaliana* |  |
| Q6NKZ8 | AtCYP714A2 | *Arabidopsis thaliana* |  |
| AF318500 | AtKAO1 | *Arabidopsis thaliana* | Kaurenoic acid oxidase (KAO) |
| AF318501 | AtKAO2 | *Arabidopsis thaliana* | Kaurenoic acid oxidase (KAO) |
| AF047719 | AtKO | *Arabidopsis thaliana* | Kaurene oxidase (KO) |
| ACQ99375 | CaKO | *Coffea arabica* | Kaurene oxidase (KO) |
| HQ658173 | CamKAO | *Castanea mollissima* |  |
| KT382342 | CfCYP71D381 | *Coleus forskohlii* | Forskolin biosynthesis |
| KT382346 | CfCYP76AH10 | *Coleus forskohlii* | Forskolin biosynthesis |
| KT382349 | CfCYP76AH11 | *Coleus forskohlii* | Forskolin biosynthesis |
| KT382358 | CfCYP76AH15 | *Coleus forskohlii* | Forskolin biosynthesis |
| KT382359 | CfCYP76AH16 | *Coleus forskohlii* | Forskolin biosynthesis |
| KT382360 | CfCYP76AH17 | *Coleus forskohlii* | Forskolin biosynthesis |
| KT382348 | CfCYP76AH8 | *Coleus forskohlii* | Forskolin biosynthesis |
| KT382347 | CfCYP76AH9 | *Coleus forskohlii* | Forskolin biosynthesis |
| NP_001267703 | CsKO | *Cucumis sativus* | Kaurene oxidase (KO) |
| AF212991 | CumKAO | *Cucurbita maxima* | Kaurenoic acid oxidase (KAO) |
| AF212990 | CumKO | *Cucurbita maxima* | Kaurene oxidase (KO) |
| KR350668 | ElCYP71D445 | *Euphorbia lathyris* | Casbene oxidase |
| KR350669 | ElCYP726A27 | *Euphorbia lathyris* | Casbene oxidase |
| KX428471 | EpCYP71D365 | *Euphorbia peplus* | Casbene oxidase |
| KJ026362 | EpCYP726A19 | *Euphorbia peplus* | Casbene synthase |
| KF986823.1 | EpCYP726A4 | *Euphorbia peplus* | Casbene oxidase |
| KF773141 | GbCYP716B | *Ginkgo biloba* | taxoid-9α-hydroxylase |
| KHN31869 | GsKO | *Glycine soja* | Kaurene oxidase (KO) |
| FR666915 | HaKAO1 | *Helianthus annuus* | Kaurenoic acid oxidase (KAO) |
| FR666916 | HaKAO2 | *Helianthus annuus* | Kaurenoic acid oxidase (KAO) |
| AF326277 | HvKAO1 | *Hordeum vulgare* | Kaurenoic acid oxidase (KAO) |
| KX060559 | JcCYP71D495 | *Jatropha curcas* | Casbene oxidase |
| KF986815 | JcCYP726A20 | *Jatropha curcas* | Casbene oxidase |
| KF986816 | JcCYP726A21 | *Jatropha curcas* |  |
| KF986818 | JcCYP726A23 | *Jatropha curcas* |  |
| KF986819 | JcCYP726A24 | *Jatropha curcas* |  |
| KX060558 | JcCYP726A35 | *Jatropha curcas* | Casbene oxidase |
| JF929910 | JcKO | *Jatropha curcas* | Kaurene oxidase (KO) |
| AB370238 | LsKAO | *Lactuca sativa* | Kaurenoic acid oxidase (KAO) |
| ADE06669 | McKO | *Momordica charantia* | Kaurene oxidase (KO) |
| KF437682 | MdKAO | *Malus domestica* | Kaurenoic acid oxidase (KAO) |
| AY563549 | MdKO | *Malus domestica* | Kaurene oxidase (KO) |
| XP_010089925 | MnKO | *Morus notabilis* | Kaurene oxidase (KO) |
| XM_013607618 | MtKAO | *Medicago truncatula* | Kaurenoic acid oxidase (KAO) |
| AF116915 | NtCYP51 | *Nicotiana tabacum* | obtusifoliol 14-alpha demethylase |
| AF166332 | NtCYP71D16 | *Nicotiana tabacum* | CBT-ol oxidase |
| Q7XHW5 | OsCYP714B1 | *Oryza sativa* |  |
| Q0DS59 | OsCYP714B2 | *Oryza sativa* |  |
| AK109526 | OsCYP714D1 | *Oryza sativa* |  |
| AK107418 | OsCYP71Z6 | *Oryza sativa* |  |
| AK070167 | OsCYP71Z7 | *Oryza sativa* | Cassadiene oxidase |
| AK059010 | OsCYP76M5 | *Oryza sativa* |  |
| AK101003 | OsCYP76M6 | *Oryza sativa* | Oryzalexin synthase |
| AK105913 | OsCYP76M7 | *Oryza sativa* |  |
| AK069701 | OsCYP76M8 | *Oryza sativa* | Oryzalexin synthase |
| AK071864 | OsCYP99A2 | *Oryza sativa* | Pimaradiene oxidase |
| AK071546 | OsCYP99A3 | *Oryza sativa* | Pimaradiene oxidase |
| Q5VRM7 | OsKAO | *Oryza sativa* | Kaurenoic acid oxidase (KAO) |
| Q5Z5R4 | OsKO2 | *Oryza sativa* | Kaurene oxidase (KO) |
| AY579214 | OsKOL4 | *Oryza sativa* | Kaurene oxidase (KO) |
| AY660664 | OsKOL5 | *Oryza sativa* | Kaurene oxidase (KO) |
| KJ845667 | PbCYP720B2 | *Pinus banksiana* | Hydroxyl abietene oxidase |
| KJ845671 | PcCYP720B1 | *Pinus contorta* | Abietadienol/abietadienal oxidase |
| KJ845675 | PcCYP720B11 | *Pinus contorta* |  |
| KJ845676 | PcCYP720B12 | *Pinus contorta* | Hydroxyl abietene oxidase |
| KJ845672 | PcCYP720B2 | *Pinus contorta* | Hydroxyl abietene oxidase |
| AEK01241 | PcKO | *Pyrus communis* | Kaurene oxidase (KO) |
| BAK19917 | PhpKO | *Physcomitrella patens* | Kaurene oxidase (KO) |
| AF537321 | PsaKAO1 | *Pisum sativum* | Kaurenoic acid oxidase (KAO) |
| AF537322 | PsaKAO2 | *Pisum sativum* | Kaurenoic acid oxidase (KAO) |
| AAP69988 | PsaKO | *Pisum sativum* | Kaurene oxidase (KO) |
| HM245408 | PsiCYP720B10 | *Picea sitchensis* |  |
| HM245397 | PsiCYP720B12 | *Picea sitchensis* | Hydroxyl abietene oxidase |
| HM245398 | PsiCYP720B15 | *Picea sitchensis* |  |
| HM245399 | PsiCYP720B16 | *Picea sitchensis* |  |
| HM245402 | PsiCYP720B2 | *Picea sitchensis* | Hydroxyl abietene oxidase |
| HM245403 | PsiCYP720B4 | *Picea sitchensis* | Diterpene C-18 oxidase |
| HM245406 | PsiCYP720B7 | *Picea sitchensis* |  |
| HM245407 | PsiCYP720B8 | *Picea sitchensis* |  |
| HM245410 | PsiCYP720B9 | *Picea sitchensis* |  |
| AY779541 | PtaCYP720B1 | *Pinus taeda* | Abietadienol/abietadienal oxidase |
| XP_006386514 | PtrKO | *Populus trichocarpa* | Kaurene oxidase (KO) |
| HM003112 | PypKO | *Pyrus pyrifolia* | Kaurene oxidase (KO) |
| KF986809 | RcCYP726A13 | *Ricinus communis* |  |
| KF986810 | RcCYP726A14 | *Ricinus communis* | Casbene oxidase |
| KF986811 | RcCYP726A15 | *Ricinus communis* | Neocembrene oxidase |
| KF986812 | RcCYP726A16 | *Ricinus communis* | Casbene oxidase |
| KF986813 | RcCYP726A17 | *Ricinus communis* | Casbene oxidase |
| KF986814 | RcCYP726A18 | *Ricinus communis* | Casbene oxidase |
| KP091843 | RoCYP76AH22 | *Rosmariuns officinalis* | Hydroxyferruginol synthase |
| KP091844 | RoCYP76AH23 | *Rosmariuns officinalis* | Hydroxyferruginol synthase |
|  | RoCYP76AH4 | *Rosmariuns officinalis* | Hydroxyferruginol synthase |
| KX431219 | RoCYP76AK7 | *Rosmariuns officinalis* | C20 oxidase |
| KX431220 | RoCYP76AK8 | *Rosmariuns officinalis* | C20 oxidase |
| U74319 | SbCYP51 | *Sorghum bicolor* | obtusifoliol 14-alpha demethylase |
| KP091842 | SfCYP76AH24 | *Salvia fruticosa* | Hydroxyferruginol synthase |
| KX431218 | SfCYP76AK6 | *Salvia fruticosa* | C20 oxidase |
| Solyc08g005650.2.1 | SlCYP71BN1 | *Solanum lycopersisum* |  |
| JX422213 | SmCYP76AH1 | *Salvia miltiorrhiza* | Ferruginol synthase |
| KR140168 | SmCYP76AH3 | *Salvia miltiorrhiza* | Hydroxyferruginol synthase |
| KR140169 | SmCYP76AK1 | *Salvia miltiorrhiza* | C20 oxidase |
| KP337688 | SmCYP76AK2 | *Salvia miltiorrhiza* |  |
| KP337689 | SmCYP76AK3 | *Salvia miltiorrhiza* |  |
| KJ606394 | SmKO | *Salvia miltiorrhiza* | Kaurene oxidase (KO) |
| KT157042 | SpCYP71BE52 | *Salvia pomifera* | C2 oxidase |
| KT157044 | SpCYP76AH24 | *Salvia pomifera* | Hydroxyferruginol synthase |
| KT157045 | SpCYP76AK6 | *Salvia pomifera* | C20 oxidase |
| AY364317 | SrKO1 | *Stevia rebaudiana* | Kaurene oxidase (KO) |
| AY995178 | SrKO2 | *Stevia rebaudiana* | Kaurene oxidase (KO) |
| ADZ55286 | TaKO | *Triticum aestivum* | Kaurene oxidase (KO) |
| AY518383 | TcaCYP725A6, T2OH | *Taxus canadensis* | taxoid-2α-hydroxylase |
| AF318211 | TcuCYP725A1, T10OH | *Taxus cuspidata* | taxoid-10β-hydroxylase |
| AY056019 | TcuCYP725A2, T13OH | *Taxus cuspidata* | taxoid-13α-hydroxylase |
| AY188177 | TcuCYP725A3, T14OH | *Taxus cuspidata* | taxoid-14β-hydroxylase |
| AY289209 | TcuCYP725A4, T5OH | *Taxus cuspidata* |  |
| AY307951 | TcuCYP725A5, T7OH | *Taxus cuspidata* | taxoid-7β-hydroxylase |
| MG696754.1 | VacCYP76BK1 | *Vitex agnus-castus* |  |
| JQ086553 | VvKO | *Vitis vinifera* | Kaurene oxidase (KO) |
| ACG38493 | ZmKO | *Zea mays* | Kaurene oxidase (KO) |
| XP_034572432.1 | SvCYP89A2-like | *Setaria viridis* | unknown function |
| Q42602.2 | AtCYP89A2 | *Arabidopsis thaliana* | unknown function |
| KAF6985951 | TaCYP89A2-like | *Triticum aestivum* | unknown function |

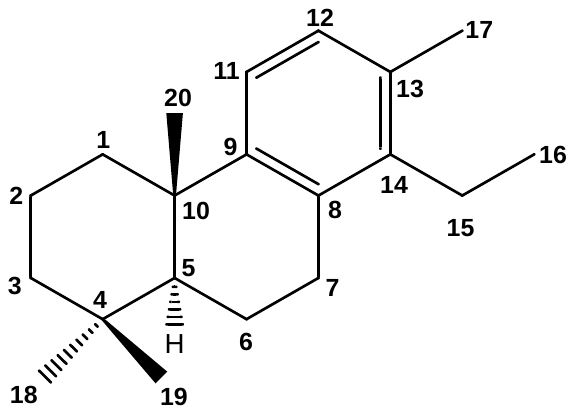
**Table S6. NMR data of hordetriene (compound 2) (C_6_D_6_)**

| **Pos.** | **^13^C: δ [ppm]** | **^1^H: δ [ppm] m (J[Hz]^a^)** | **HMBC (H 🡪C)** | **ROE^d^** |
| --- | --- | --- | --- | --- |
| 1 | 39.6 | 2.206 m β; 1.344 td-like (13.2/3.8) α |  | 11,20; |
| 2 | 19.8 | 1.670 qt-like (13.9/3.4) β;  1.5054 m α | 1 4,10 | 20; |
| 3 | 41.9 | 1.409 m β;  1.138 td-like (13.5/4.0) α | 4 |  |
| 4 | 33.4 | --- |  |  |
| 5 | 50.0 | 1.243 dd (12.6/2.2) α |  | 18 |
| 6 | 19.7 | 1.797 ddt-like (13.2/7,8/1.9) α; 1.613 m β |  |  |
| 7 | 28.0 | 2.849 ddd (17.1/6.6/1.3) β; 2.643 ddd (17.1/11.6/7.8) α |  |  |
| 8 | 132.5^b^ | --- |  |  |
| 9 | 148.5 | --- |  |  |
| 10 | 38.1 | --- |  |  |
| 11 | 122.4 | 7.092 d (8.0) |  |  |
| 12 | 128.4^c^ | 7.020 d (8.0) |  | 17 |
| 13 | 132.6^b^ | --- |  |  |
| 14 | 140.2 | --- |  |  |
| 15 | 22.4 | 2.52 m | 16 |  |
| 16 | 13.4 | 1.035 t (7.5) | 14,15 |  |
| 17 | 19.5 | 2.209 s | 12,13,14 |  |
| 18 | 33.4 | 0.923 s | 3,4,5,19 |  |
| 19 | 21.8 | 0.899 s | 3,4,5,18 | 20 |
| 20 | 25.2 | 1.197 br s | 1,5,9,10 |  |

^a^ “-like” multiplicities: the given value represents the distance of the multiplet lines, not the exact coupling constant; ^b^ might be interchanged; ^c^ ^13^C chemical shift of HSQC correlation peak; ^d^ each NOE is listed only once.

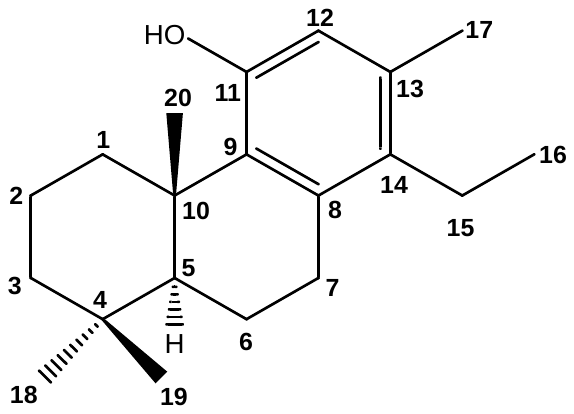
**Table S7. NMR data of 11-hydroxy-hordetriene**

**(compound 3) (C_6_D_6_)**

| **Pos.** | **^13^C: δ [ppm]** | **^1^H: δ [ppm] m (J[Hz]^a^)** | **HMBC (H 🡪 C)** | **ROE (H ⭤ H)^b^** |
| --- | --- | --- | --- | --- |
| 1 | 37.26 | 3.848 br dt (13.4/3.7) β; 1.350 td (13.4/3.7) α | 2,3,5,10; 2,10 | 20; |
| 2 | 19.98 | 1.824 dtt (14.0/13.4/3.7) β; 1.582 dt-like (14.0/3.7) α | 1,3,4,10; 4,10 | 20; |
| 3 | 41.83 | 1.453 dtd-like (13.1/3.5/1.5) β; 1.234 td-like (13.4/4.2) α | 2,4; 2,4,18,19 |  |
| 4 | 33.77 | --- |  |  |
| 5 | 52.88 | 1.276 dd (12.2/1.3) α | 4,6,10,18,20 | 18 |
| 6 | 19.64 | 1.769 br dd-like (12.9/6.8) α; 1.521 qt-like (12.5/5.4) β | 8,10; 5,10 | 18,19; 19 |
| 7 | 30.64 | 2.791 ddd (16.8/5.4/1.5) β; 2.614 ddd (16.8/12.6/6.8) α | 5,6,8,9,14; 6,8 | 15; |
| 8 | 136.02 | --- |  |  |
| 9 | 133.62 | --- |  |  |
| 10 | 39.72 | --- |  |  |
| 11 | 152.11 | --- |  |  |
| 12 | 116.67 | 5.668 br s | 9,11,14,17 | 11-OH,17 |
| 13 | 133.29 | --- |  |  |
| 14 | 132.84 | --- |  |  |
| 15 | 21.99 | 2.475 q (7.5) | 8,13,14,16 | 17 |
| 16 | 13.74 | 1.030 t (7.5) | 14,15 |  |
| 17 | 19.12 | 2.097 br s | 12,13,14 |  |
| 18 | 33.88 | 0.960 s | 3,4,5,19 |  |
| 19 | 22.36 | 0.954 s | 3,4,5,18 | 20 |
| 20 | 20.05 | 1.549 s | 1,5,9,10 |  |
| 11-OH | --- | 3.726 s | 9,11,12 |  |

^a^ “-like” multiplicities: the given value represents the distance of the multiplet lines, not the exact coupling constant; ^b^ each NOE is listed only once.

**Table S8. Oligonucleotides used in this study.**

| **Gene** | **Application** | **Direction** | **Sequence (5’ 🡪 3’)** |
| --- | --- | --- | --- |
| GenBank accession no. M60175 (HvUBIQUITIN) | qPCR | Forward | ACCCTCGCCGACTACAACAT |
|  |  | Reverse | CAGTAGTGGCGGTCGAAGTG |
| HORVU2Hr1G004540 (HvKSL4) | qPCR | Forward | GTTATCTCTGCGCTGCTGCC |
|  |  | Reverse | GAGGATCCTCCCATTTCTCAGC |
|  | truncate transit peptide | Forward | TTGGTCTCAACATAATGGCTTACGTTGAATCTAGACC |
|  |  | Reverse | TTGGTCTCAACAAACCAATTCGTTTTGAGACAAAATG |
| HORVU2Hr1G004620 (HvCPS2) | qPCR | Forward | GCGTCTGCAGCCCATGAGA |
|  |  | Reverse | GCGGTCTTCCTCCCTCTGC |
|  | truncate transit peptide | Forward | TTGGTCTCAACATAATGGTTTTGTCCTCTAAATCTCCA |
|  |  | Reverse | TTGGTCTCAACAAGGGTTAACTTCGTAACCATGTTG |
| HORVU2Hr1G004550 (HvCYP89E31) | qPCR | Forward | GCGTCTGCAGCCCATGAGA |
|  |  | Reverse | GCGGTCTTCCTCCCTCTGC |
| HORVU2Hr1G004530 (HvCYP99A67) | qPCR | Forward | GCGATCATATCGGATATGTTCACG |
|  |  | Reverse | TCTTGTTGTCAAAGGTACGACGC |
| HORVU2Hr1G004480 (HvCYP99A66) | qPCR | Forward | GTTCGTCGACGCCCTCACT |
|  |  | Reverse | CCGTGAACATATCCAATATGATCG |
